# Longitudinal proteomic module profiles vary across human monocyte differentiation and polarization conditions

**DOI:** 10.64898/2026.06.24.734366

**Authors:** Roberto Navarro Quiroz, Yiris Diaz-Olmos, Noelia Geribaldi-Dóldan, Cecilia Fernández-Ponce, Eloina Zarate Peñata, Katherine Escorcia Lindo, Ibeth Karina Luna-Rodríguez, Andrea Jaruffe Pinilla, Yesit Bello Lemus, Lisandro Pacheco Lugo, Leonardo Pacheco Londoño, Antonio Acosta-Hoyos, Nataly Galan Freyle, Elkin Navarro Quiroz

**Affiliations:** Center for Research in Critical Dynamics, Barranquilla, Colombia; Tecnológico de Antioquia—Institución Universitaria, Medellín, Colombia; Facultad de Ciencias Básicas y Biomédicas, Centro de Investigaciones en Ciencias de la Vida (CICV), Universidad Simón Bolívar, Barranquilla, Colombia; División de Ciencias de la Salud, Programa de Medicina, Universidad del Norte, Barranquilla, Colombia; Instituto de Investigación e Innovación Biomédica de Cádiz (INiBICA), Cádiz, Spain; Departamento de Anatomía, Facultad de Medicina, Universidad de Cádiz, Cádiz, Spain; Departamento de Biomedicina, Biotecnología y Salud Pública, Facultad de Medicina, Universidad de Cádiz, Cádiz, Spain

**Keywords:** monocytes, macrophage polarization, dendritic cells, osteoclasts, proteomics, longitudinal analysis, immunometabolism, protein modules

## Abstract

Human monocyte differentiation involves changes in multiple protein programs, but comparisons across culture conditions can conflate source variation with differentiation time. We reanalyzed public proteomic measurements arranged in four source blocks across M1-polarizing, M2-polarizing, dendritic-cell and osteoclast conditions at days 2, 4, 6, 8 and 10. The original experiment used three individual-donor preparations and one preparation pooled from 40 donors; these are not four individual donors. Eight predefined protein-module scores formed an 80-observation matrix. A multivariate model tested joint condition and condition-by-time terms beyond source block and categorical time, using 9,999 permutations that preserved complete block-specific culture trajectories. The joint condition terms accounted for an additional 31.37% of total module-score variation (pseudo-F = 4.257567; permutation p = 0.0002). The association persisted after omission of each source block, with incremental R^2^ of 0.315–0.390. Secondary day-specific tests detected differences at days 4–10, whereas day 2 did not meet the false-discovery criterion. Descriptive trajectories showed a shared increase in glycolysis/pentose-phosphate scores alongside condition-dependent mitochondrial, redox and proteostasis profiles. At day 10, M2-labelled cultures had the lowest mean mitochondrial score, whereas M1-labelled cultures had the highest three-protein resolution-associated score. All eight module tests survived multiplicity correction, and no single-module omission abolished the multivariate association. These findings describe condition-associated configurations within four heterogeneous source preparations. Equal weighting of source blocks does not estimate a mean across individual donors. The analysis does not establish pathway activity, single-cell trajectories, population-level replication or external biological validation.

## 1 Introduction

Monocytes can acquire different molecular and cellular properties in response to differentiation and activation signals. Human macrophage activation extends beyond a binary M1/M2 classification, as illustrated by experimental transcriptome profiles spanning multiple stimuli (1). Consequently, a culture label provides information about how cells were generated, but does not by itself establish every function conventionally associated with that label. This distinction becomes particularly relevant when comparing macrophage-polarizing conditions with monocyte-derived dendritic-cell and osteoclast differentiation, which represent different biological processes (2) (3).

The timing of observation also matters. An early culture and its later counterpart may differ because of processes shared across differentiation conditions, because of condition-dependent remodeling, or both. Pooling observations from different days can obscure those components and inflate the apparent number of independent biological replicates. Serial cultures permit description of temporal profiles while preserving their source pairing. However, a source series generated from pooled cells is not equivalent to an individually resolved donor. Such bulk measurements characterize source-matched cultures over time; they do not follow the same individual cells or establish lineage relationships.

Proteomic measurements complement transcriptomic descriptions by quantifying proteins involved in metabolism, signaling, lipid handling and protein maintenance. Summarizing a prespecified set of proteins can make a broad comparison easier to interpret, but a module score remains dependent on its membership, coverage, sign convention and aggregation rule. A signed redox score, for example, can combine proteins involved in oxidant production with antioxidant proteins assigned an opposite sign. It therefore summarizes the balance within that measured panel rather than the amount of reactive oxygen species. Partial coverage poses an additional limitation: a score based on LRP1, APOE and TGFB1 cannot substitute for an efferocytosis assay.

Primary human macrophage studies have reported activation-associated metabolic profiles (4), differences among M2-associated metabolite profiles (5), and lipid remodeling during the inflammatory response and its resolution (6). These findings motivate examination of metabolic and related protein panels. They also make it necessary to distinguish evidence from different measurement layers. A protein-abundance pattern and a metabolite-abundance pattern measured in unrelated cultures do not constitute paired multiomic evidence.

Here we asked whether eight predefined proteomic modules retain an association with human monocyte-derived differentiation and polarization conditions after accounting for source block and differentiation day. We used PXD016245, a serial proteomic resource with four source blocks, four conditions and five post-baseline time points (7). The source experiment includes individual and pooled preparations (3). The primary comparison tested the additional condition and condition-by-time terms jointly. We then described the observed profiles, the temporal distribution of the association and its sensitivity to omission of source blocks, module membership and scaling. Our objective was to establish what this small, structured proteomic dataset supports about condition-associated configurations, without equating module abundance with function or source blocks with four individual donors.

## 2 Materials and methods

### 2.1 Study design and analysis history

This was a retrospective analysis of public omics data. The time-adjusted analysis plan was locked on 19 June 2026 before execution of that specific analysis. Earlier descriptive and pooled analyses had already examined the same dataset. Thus, prespecification refers to the revised models, permutation scheme, primary criterion and sensitivity analyses; it does not imply prospective recruitment, a public clinical-study registration or complete absence of prior knowledge of the data. The archived plan, input and permutation assignments accompany an independent primary-result verifier in Supplementary Data 3; the frozen secondary statistics are provided in Supplementary Data 1. The editorial revision adds descriptive summaries of the frozen scores, including condition-by-day means, source block ranges and paired day-10-minus-day-2 changes. These summaries were not new prespecified hypothesis tests.

### 2.2 Proteomic source and sample structure

PXD016245 was obtained from PRIDE/ProteomeXchange. The monocyte-differentiation chapter of Béguin’s thesis describes three individual-donor preparations and one preparation pooled from 40 donors (3, p. 94), and identifies the deposit as PXD016245 (3, p. 95; 7). The reused sample names contain prefixes A, B, C and P under M1-polarizing, M2-polarizing, dendritic-cell and osteoclast differentiation conditions. We call these four longitudinal source blocks. The historical parser stored these prefixes in a column named donor_id without an independent sample sheet. The literal correspondence of P to the pooled preparation was not established from the inspected metadata, so no inference depends on asserting that mapping.

Each source-block–condition combination was sampled at days 2, 4, 6, 8 and 10. The analytical matrix therefore contains 80 serial bulk observations, comprising 16 block-condition trajectories. Four additional day-0 monocyte observations contributed to the historical standardization but were excluded from the condition comparison because they do not form a four-condition baseline factorial. There is one observation per block-condition-day combination. Time was recovered from the original field named replicate and represented explicitly as differentiation day. The 80 measurements and 16 trajectories are not independent donor replicates; the pooled preparation likewise does not yield 40 separately measured donor outcomes. Operational culture labels preserve the source design and are not independently validated functional phenotypes.

The source-design interpretation was corrected during final editorial verification on 7 September 2026. The earlier description of four independent donors was an unsupported interpretation of filename prefixes. The correction changes the biological claim and variable description, while retaining the frozen observations, models and permutation assignments. The pooled preparation’s contributors and possible overlap with the individual preparations were not resolved; independence across four individual donors is not claimed.

### 2.3 Original proteomic processing and module construction

The dataset record identifies an Orbitrap Fusion instrument. We reused the deposited MaxQuant protein-group LFQ matrix and did not reprocess raw spectra. The archived search-parameter file specifies MaxQuant 1.5.3.30, peptide- and protein-level false-discovery thresholds of 1%, match-between-runs enabled and LFQ normalization enabled. The search FASTA is recorded as HUMAN170307.fasta. Its exact database release and sequence count were not established from the available parameter file. Supplementary Methods distinguish these original search settings from the subsequent transformations used to construct the module scores.

The historical import excluded protein groups flagged as reverse identifications, potential contaminants or identified only by site. Their overlapping union removed 346 of 6,375 rows, leaving 6,029 protein-group features across 84 LFQ sample columns. Positive LFQ values were log2-transformed; non-numeric, zero and negative values were treated as missing. Each assay feature was standardized using its observed values across all 84 samples, including the four day-0 monocytes, and the sample standard deviation. Features with more than 50% missing values or zero variance were excluded from module scoring. There was no imputation or additional sample normalization at this scoring stage. Thus, the absence of additional normalization in the reanalysis should not be confused with the source’s LFQ processing.

The eight modules covered glycolysis/pentose-phosphate proteins, mitochondrial proteins, lipid handling, redox-related proteins, inflammasome/cytokine-related proteins, resolution-associated proteins, monocyte/macrophage differentiation-associated proteins and proteostasis-associated proteins. Module membership and directions were inherited from the frozen definitions. A sample’s score was the mean of its available oriented assay-feature z-scores, requiring at least two usable features. Features assigned a negative direction were sign-reversed before averaging. Multiple eligible features assigned to the same named protein were retained: G6PD and PDIA3 each contributed two features. The scores therefore do not give equal weight to every unique protein name.

The final representation comprised 54 distinct eligible assay features and 55 feature-module memberships. APOE occurred in the lipid-handling and resolution-associated panels with opposite orientations. Expected protein coverage and eligible feature counts are distinguished in Table 1; the full mapping and missingness records are supplied in Supplementary Data 1. Because only LRP1, APOE and TGFB1 contributed to the resolution-associated panel, we refer to it as the partial resolution-associated score. No rescoring, imputation, feature replacement or change in module membership was performed for this revision.

**Table 1.** Coverage and assay-feature membership of the eight protein panels.

| Protein panel | Eligible / expected proteins | Eligible features | Interpretation constraint |
| --- | --- | --- | --- |
| Glycolysis and PPP | 8 / 10 | 9 | Two G6PD features retained |
| Mitochondrial | 10 / 11 | 10 | Protein abundance, not respiration |
| Lipid handling | 6 / 9 | 6 | APOE shared, negatively oriented |
| Redox | 8 / 9 | 8 | Positive and negative orientations |
| Inflammasome and cytokine related | 4 / 9 | 4 | Incomplete panel; not cytokine output |
| Partial resolution associated | 3 / 6 | 3 | LRP1, APOE, TGFB1 only |
| Differentiation associated | 7 / 12 | 7 | Operational abundance summary |
| Proteostasis associated | 7 / 10 | 8 | Two PDIA3 features retained |

PPP, pentose phosphate pathway. Coverage counts unique expected protein identities represented by at least one eligible feature; the score averages assay features. Across panels, 55 memberships represent 54 distinct assay-feature identifiers. Full identifiers, signs and missingness are in Supplementary Data 1.

### 2.4 Primary multivariate model and permutation inference

The response matrix contained 80 observations and eight frozen module scores. We compared the reduced model, Y ~ source_block + factor(day), with the full model, Y ~ source_block + factor(day) + condition + factor(day):condition. Least-squares fits used the squared Frobenius norm of residuals, equivalent to a Euclidean distance-based multivariate partition (8). The reduced and full design matrices had ranks 8 and 23. The tested joint condition block therefore had 15 degrees of freedom, with 57 residual degrees of freedom.

The pseudo-F statistic was (SS block/15)/(SSE full/57). Incremental R^2^ was (SSE reduced − SSE full)/SS total, measuring the additional fraction of total module-score variation accounted for by condition and its interaction with day beyond source block and shared day effects. This denominator differs from partial R^2^, and the statistic is not a condition main-effect estimate. The prespecified support criterion required permutation p ≤ 0.05 and incremental R^2^ > 0.02; the latter was a study-specific descriptive relevance criterion.

Within each source block, the four condition labels were permuted once and the same assignment applied at every day. Observations were never transferred between source blocks and day labels were not permuted. This preserves complete block-condition trajectories under the null of exchangeable condition labels within source block. From the 24^4^ = 331,776 possible assignments, 9,999 distinct non-identity assignments were sampled without replacement using NumPy PCG64 and seed 20260615. We calculated p = (b + 1)/(9,999 + 1), counting permuted statistics at least as large as the observed value with a numerical tolerance of 10^−9^. This is a Monte Carlo permutation test rather than exhaustive enumeration. Interpretation depends on the assumed exchangeability and cannot distinguish a condition-linked technical effect from a condition-linked biological effect.

### 2.5 Secondary analyses and sensitivity checks

Four leave-one-source-block-out analyses repeated the primary model using 60 observations from three retained blocks, with 15 tested-term and 38 residual degrees of freedom and 9,999 trajectory-preserving permutations. These overlapping subsets assess dependence on a source series; they do not supply independent validation cohorts or a donor-level sampling distribution. The analysis excluding P retains the original frozen scores, whose feature standardization used all 84 source observations. It is therefore not a new preprocessing analysis without pooled material. No donor-bootstrap confidence intervals were generated.

The condition-by-day interaction was examined as a secondary Freedman–Lane residual-permutation sensitivity (9). Residuals from an additive model containing source block, day and condition were permuted as complete trajectories within source block, added to the additive fitted values, and tested by comparing additive and full models. The additive design rank was 11. This sensitivity used 9,999 permutations and did not determine the primary conclusion. Finite-sample exactness was not established.

At each day, a block-adjusted condition model used 16 observations and within-source block permutations. The five p-values formed one Benjamini–Hochberg correction family (10). Each module was also tested with the primary reduced/full model; those eight p-values formed a separate correction family. We report adjusted q-values, the within-module or within-day incremental R^2^ and the test statistic. Neither family provides pairwise condition tests or establishes a precise onset of divergence.

Leave-one-module-out analyses omitted each module in turn. The reported change is incremental R^2^ with all modules minus incremental R^2^ without that module. Because omission changes both the response representation and its total-variation denominator, this quantity is a sensitivity measure rather than a causal contribution. A separate sensitivity centered and scaled each module to unit sample variance across the 80 observations. Restricted multivariate dispersion testing and a two-protein LRP1/TGFB1 sensitivity excluding APOE were not implemented in the locked analysis. These omissions remain limitations; variance scaling does not test equality of dispersion among conditions (11).

### 2.6 Audit of the archived metabolomic comparison

An earlier analysis of ST001835 (12) used three metabolite scores and 9,999 unrestricted group-label permutations, without donor or time adjustment. Audit identified chemically nonspecific metabolite-name assignments and repeated donor/time annotations not accommodated by the permutations. Its historical numerical outputs are retained solely for traceability in Supplementary Methods and Supplementary Data 1 and are withdrawn from biological interpretation. No replacement metabolomic test was performed. The defects concern that secondary branch; all used PXD016245 protein memberships were separately verified as exact gene-symbol matches.

### 2.7 Reproducibility and descriptive display

The original execution used Python 3.10.20, NumPy 2.2.6, pandas 2.3.3 and SciPy 1.15.3. An independent historical implementation reproduced the stored analysis. For the present editorial revision, a separate numerical audit again reproduced the observed statistic and all 9,999 archived primary permutation statistics. The frozen 80-row input is identified by SHA-256 1fb757edd9bbb8e2354547ab733fc04a7888cad661b19e488a6e13c4664a8e66. Supplementary Data 3 contains an independent verifier of the primary endpoint from frozen scores, selected historical source files, the plan, software requirements and reproduction instructions. It does not reconstruct scores or rerun secondary analyses.

Figures display the original scores and frozen inferential results. Condition-by-day means give equal weight to the four source blocks; ranges describe their observed spread. This is a preparation-level summary and not an individual-donor population mean, because one original preparation pooled 40 donors. Paired terminal-minus-early differences are calculated within block and condition. Scores and differences are dimensionless averages on the original feature-standardization scale, not fold changes. All 160 module-condition-day summaries and 32 terminal-change summaries are provided to make the selected descriptive examples auditable. No pairwise significance or confidence interval is inferred from visual separation.

## 3 Results

### 3.1 Complete serial sampling supports a block-adjusted comparison of partially observed modules

All 80 expected block-condition-day combinations were present, without duplicated combinations or missing module scores. Thus, each of the eight modules contributed 80 values, although the proteins underlying those scores were incompletely observed. The 16 trajectories arose from four source preparations; the source design comprises three individually obtained preparations and one pooled preparation. Neither serial measurements nor members contributing to the pool increase the number of separately resolved biological replicates. Figure 1 summarizes this structure and distinguishes the four baseline monocyte measurements from the post-baseline comparisons.

**Figure 1.**
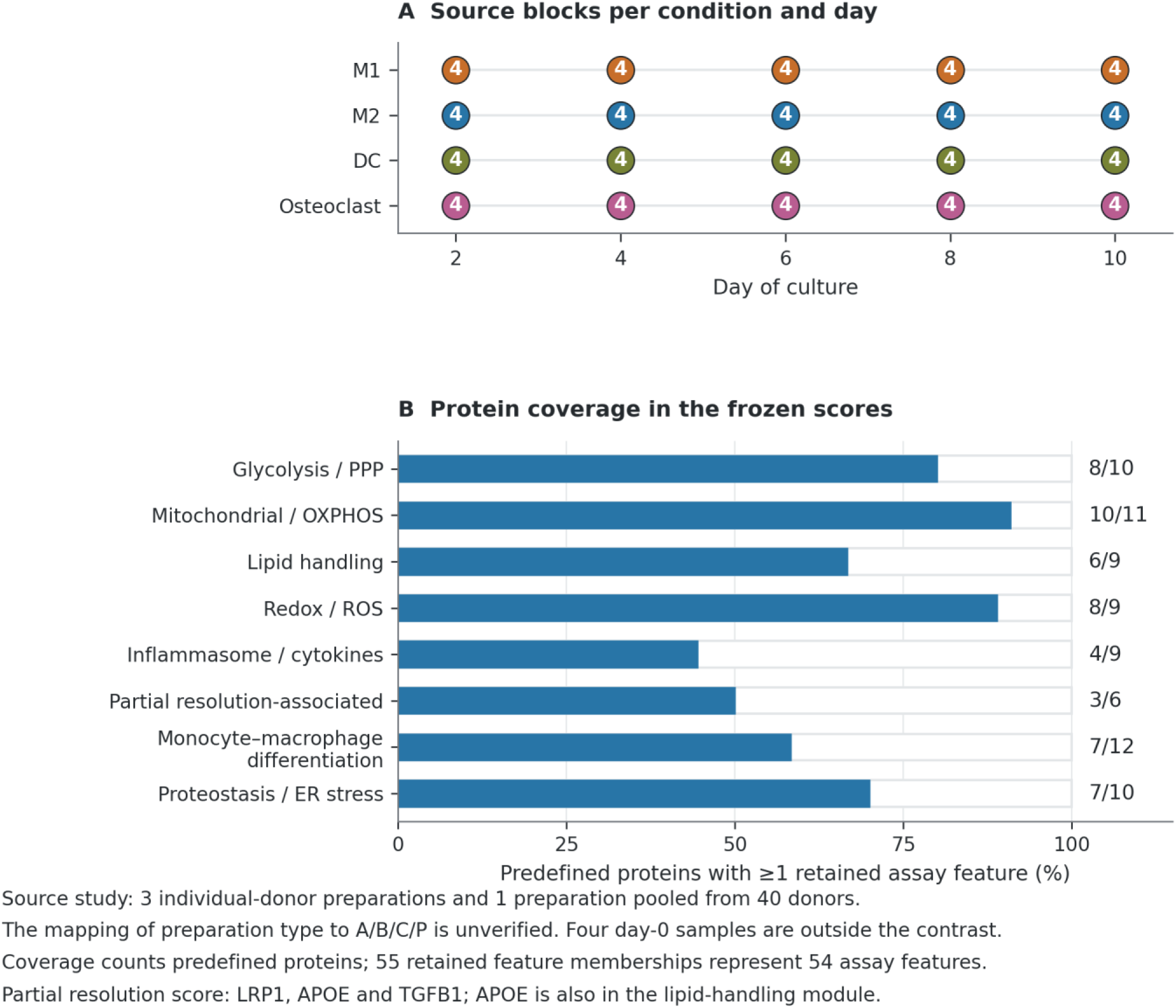
Study structure and protein-module coverage in PXD016245. (A) The longitudinal analytical matrix contains 80 observations: four source blocks (A, B, C and P), each represented in four culture conditions at days 2, 4, 6, 8 and 10. Numbers in symbols indicate the four source blocks contributing to each condition–day combination. The original study describes three individual-donor preparations and one preparation pooled from 40 donors. The mapping of these preparation types to the identifiers A/B/C/P has not been independently established; no identifier is treated as a confirmed pooled sample. DC denotes the dendritic-cell differentiation condition. M1, M2, DC and osteoclast denote the source study’s operational culture conditions, not equivalent classes of activation state. Four additional day-0 monocyte samples contributed to the original 84-sample standardization but are excluded from the longitudinal contrast. (B) Bars show the fraction of predefined protein identities represented by at least one assay feature retained in the frozen score; labels give eligible/expected protein counts. These counts differ from assay-feature counts because multiple measured features can map to one gene. The eight scores contain 55 retained feature–module memberships representing 54 unique assay-feature identifiers. The partial resolution-associated score contains LRP1, APOE and TGFB1; APOE also contributes to lipid handling. Coverage describes what was measured and retained, not functional completeness. Figure source data are provided in Supplementary Data 2.

Protein coverage ranged from 4 of 9 expected inflammasome/cytokine-related proteins to 10 of 11 mitochondrial proteins. Coverage was 8/10 for glycolysis/pentose-phosphate, 6/9 for lipid handling, 8/9 for redox, 3/6 for the partial resolution panel, 7/12 for differentiation-associated proteins and 7/10 for proteostasis-associated proteins (Table 1). Eligible assay-feature counts ranged from three to ten per module. These differences mean that a complete module-score matrix does not represent complete pathway coverage. In particular, MERTK, AXL and IL10 were absent from the partial resolution panel, and XBP1, ATF4 and DDIT3 were absent from the proteostasis panel. Interpretations below therefore refer to the observed protein panels.

### 3.2 Shared early-to-terminal changes coexist with different terminal protein profiles

The block-level trajectories provide a biological description of the adjusted multivariate result (Figure 2; Supplementary Data 2). Glycolysis/pentose-phosphate scores increased between days 2 and 10 in every block-condition pair, representing 16 paired changes across four source blocks. Condition means changed from −0.847 to 0.364 in M1, −1.343 to 0.224 in M2, −0.865 to 0.287 in dendritic-cell cultures, and −0.826 to 1.168 in osteoclast cultures. The mean paired change in osteoclast cultures was 1.994 score units, with source block changes ranging from 1.686 to 2.385. This shared direction, combined with different magnitudes, illustrates why both day and condition belong in the comparison. It does not measure glycolytic flux.

**Figure 2.**
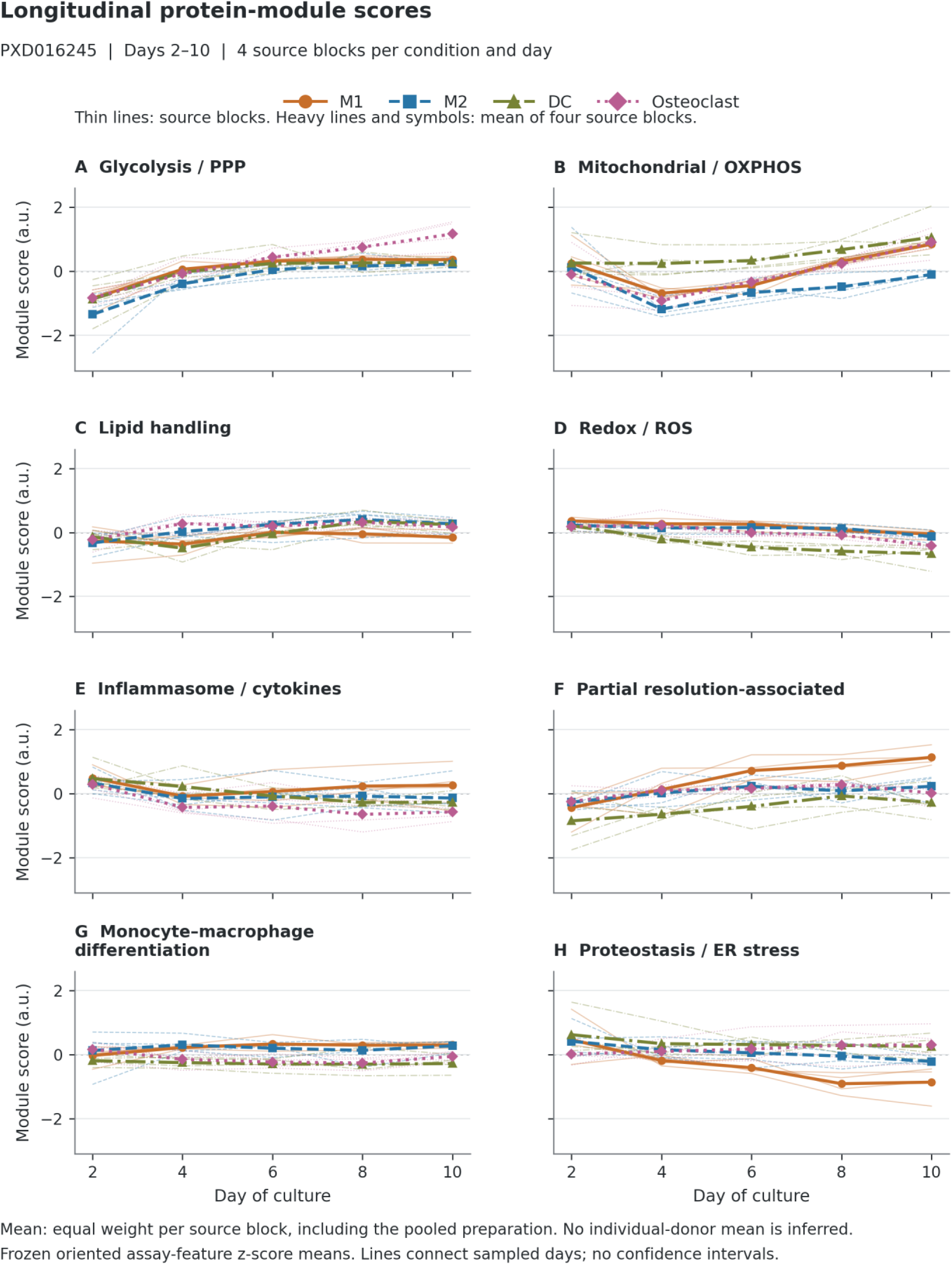
Longitudinal profiles of the eight frozen protein-module scores. Panels A–H show the predefined modules for the 80 PXD016245 longitudinal observations. Thin lines connect the five sampled days for each source block within a culture condition; heavy lines and symbols show the unweighted mean of four source blocks at each condition and day. Each block has equal weight in this descriptive mean, including the pooled preparation; it is not an average across four individual donors or a donor-weighted estimate across the pool constituents. Color, line style and symbol distinguish the four conditions. All panels use the same vertical scale. Values are the frozen means of available, prespecified-sign-oriented assay-feature z-scores, with feature standardization inherited from all 84 samples, including day-0 monocytes. These are descriptive, unadjusted scores; model adjustment is applied in the analyses in Figures 3 and 4. Connecting segments aid reading between sampled days and do not estimate a continuous trajectory. Individual lines show observed source-block variation, not confidence intervals. The partial resolution-associated score is limited to LRP1, APOE and TGFB1. Module names do not imply measured pathway activity, flux or function. a.u., arbitrary score units; DC, dendritic-cell differentiation condition; PPP, pentose phosphate pathway; OXPHOS, oxidative phosphorylation; ROS, reactive oxygen species; ER, endoplasmic reticulum. Figure source data are provided in Supplementary Data 2.

The mitochondrial panel showed a different condition ordering. At day 10, its mean score was −0.099 in M2, compared with 0.846 in M1, 1.065 in dendritic-cell cultures and 0.906 in osteoclast cultures. All four M2 scores, ranging from −0.194 to 0.057, were below the day-10 scores of the other conditions. However, block-specific changes from day 2 to day 10 were heterogeneous: three of four increased in each of M1, dendritic-cell and osteoclast cultures, and two of four increased in M2. The terminal contrast therefore should not be restated as a uniform mitochondrial decline in M2 or as a measurement of respiratory capacity.

Redox scores decreased between days 2 and 10 in all 16 block-condition pairs. Mean scores changed from 0.359 to −0.050 in M1, 0.228 to −0.126 in M2, 0.210 to −0.664 in dendritic-cell cultures and 0.188 to −0.410 in osteoclast cultures. The largest mean decrease was in dendritic-cell cultures, −0.875 units, although its source block changes ranged from −1.427 to −0.430. This panel combines positively oriented CYBB, NCF2 and NCF4 with negatively oriented SOD2, GPX1, PRDX1, PRDX3 and CAT. Lower scores consequently indicate a shift within that signed abundance panel, not directly observed lower oxidant production.

Proteostasis-associated scores decreased in every source block of the M1 and M2 conditions. Their means changed from 0.459 to −0.865 and from 0.411 to −0.217, respectively. The day-10 means in dendritic-cell and osteoclast cultures were 0.251 and 0.305. Individual terminal scores nevertheless crossed zero in both latter conditions, and one osteoclast source block showed a negative terminal-minus-early change. The pattern describes relative chaperone, protein-folding and related protein abundance; the missing stress-response components prevent its interpretation as direct evidence of endoplasmic-reticulum stress.

The partial resolution-associated score further illustrates the limits of conventional polarization labels. Its M1 mean increased from −0.442 at day 2 to 1.128 at day 10, with increases in all four source blocks. Terminal M1 scores ranged from 0.884 to 1.520, above every terminal source block score in the other conditions; their corresponding means were 0.221 in M2, −0.277 in dendritic-cell cultures and 0.013 in osteoclast cultures. This result concerns LRP1, APOE and TGFB1 abundances. It does not demonstrate that M1-labelled cultures perform more efferocytosis or resolve inflammation more effectively. The reported orderings are descriptive observations, with complete source block ranges in Supplementary Data 2, and were not subjected to new pairwise hypothesis tests.

### 3.3 Condition and its temporal component account for variation beyond source block and shared time

The joint condition block accounted for 31.37% of total multivariate module-score variation after source block and shared day effects were included (incremental R^2^ = 0.313663; pseudo-F = 4.257567; permutation p = 0.0002; Figure 3A). Both prespecified criteria were met. The block-plus-day model accounted for 40.64% of total variation and the full model for 72.00%; these are descriptive decompositions of the same fitted models rather than additional tests. The remaining full-model residual fraction was 28.00%.

**Figure 3.**
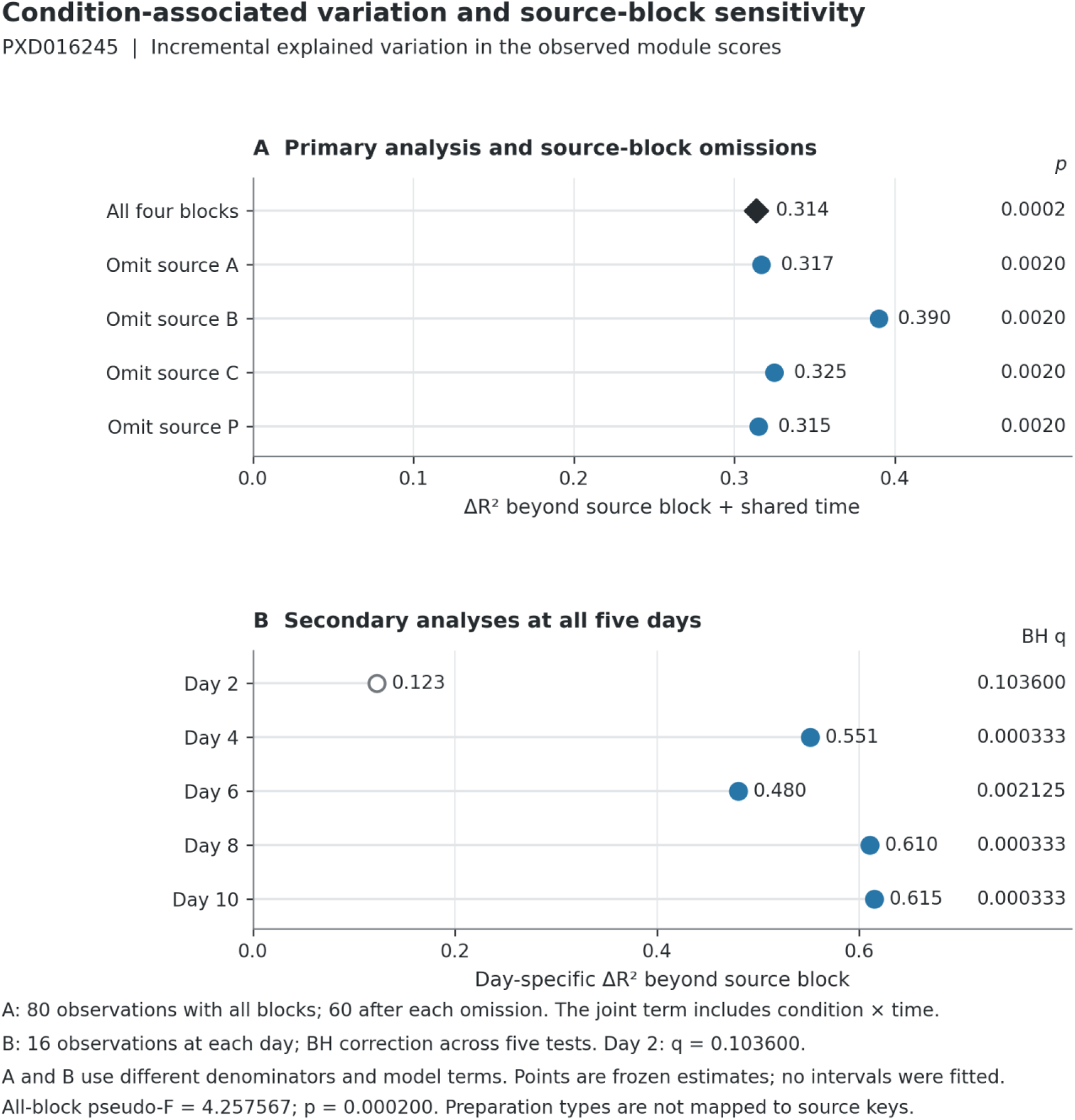
Condition-associated variation after source-block/time adjustment and sensitivity to source-block omission. (A) Points show the frozen incremental coefficient of determination, ΔR^2^ = (SSE_reduced − SSE_full)/SST, for the joint condition-associated block beyond source block and shared categorical time. The full model adds condition and condition-by-time terms. The all-block estimate (diamond; 80 observations) is 0.313663, with pseudo-F = 4.257567 and permutation p = 0.000200. Circles show estimates after separately omitting each source block (60 observations per analysis); the corresponding frozen permutation p values are printed at right. “Omit source P” denotes removal of the block labelled P; it is not presented as a confirmed pool-exclusion analysis. (B) Secondary source-block-adjusted analyses performed separately at all five sampled days, with 16 observations per day. Benjamini–Hochberg q values refer to the family of five day-specific tests. Day 2 is shown with an open symbol (q = 0.103600); this does not establish equivalence between conditions. The per-day analyses and the longitudinal joint-block analyses have different model terms and total-variation denominators. Points are frozen estimates; no confidence intervals were fitted, and source-block omissions do not constitute independent replication. Figure source data are provided in Supplementary Data 2.

This result concerns the combined condition and condition-by-day terms. It establishes an adjusted association with the culture labels within the observed dataset, but does not partition the 31.37% into independently tested condition and interaction effects. In the archived null distribution, one of 9,999 sampled assignments produced a statistic at least as large as the observed value. The conclusion was therefore supported by the structured permutation comparison rather than inferred from unadjusted visual clustering.

### 3.4 Later sampling days show adjusted differences without a monotonic temporal progression

The secondary day-specific analyses localized the multivariate association across the sampling schedule (Figure 3B; Table 2). At day 2, incremental R^2^ was 0.122569, with pseudo-F = 1.503224 and q = 0.103600. At days 4, 6, 8 and 10, incremental R^2^ values were 0.551451, 0.480102, 0.610152 and 0.614912, respectively, and all four tests survived correction across the five days. The corresponding q-values were 0.000333, 0.002125, 0.000333 and 0.000333.

**Table 2.** Secondary day-specific condition associations after source block adjustment.

| Day | Observations | Incremental R<sup>2</sup> | Pseudo-F | p-value | BH q-value |
| --- | --- | --- | --- | --- | --- |
| 2 | 16 | 0.122569 | 1.503224 | 0.103600 | 0.103600 |
| 4 | 16 | 0.551451 | 7.542594 | 0.000200 | 0.000333 |
| 6 | 16 | 0.480102 | 5.065515 | 0.001700 | 0.002125 |
| 8 | 16 | 0.610152 | 8.872380 | 0.000200 | 0.000333 |
| 10 | 16 | 0.614912 | 7.615108 | 0.000200 | 0.000333 |

Each day includes four source blocks under four conditions; degrees of freedom for condition and residuals are 3 and 9. q-values use one five-test Benjamini–Hochberg family. The incremental R<sup>2</sup> denominator is total variation within that day. These secondary tests do not establish equivalence at day 2 or a precise onset of differentiation.

The estimates did not increase monotonically: the fraction attributed to condition declined between days 4 and 6 before rising at later times. Each estimate uses the total variation within its own day and therefore does not measure an absolute amount of biological divergence on a common variance denominator. The secondary interaction sensitivity was compatible with condition-dependent temporal patterns (pseudo-F = 1.613093; incremental R^2^ = 0.095072; p = 0.0004). Together, these analyses support time-dependent differences in the observed module configurations. They do not identify day 4 as a precise onset, establish equivalence at day 2, or make differences between day-specific significance levels themselves an interaction test.

### 3.5 The association persists across source block omissions and module sensitivities

All four leave-one-source-block-out analyses retained the prespecified joint criterion (Figure 3A). Incremental R^2^ ranged from 0.315012 after omitting source P to 0.389790 after omitting source B; omission of A and C gave 0.316772 and 0.324852. The corresponding pseudo-F statistics ranged from 2.946285 to 5.089541, and all four permutation p-values were 0.002000. Specifically, excluding P left 60 observations with pseudo-F = 2.946285 and incremental R^2^ = 0.315012. These overlapping analyses show that removal of any one source series did not abolish the conditional association. They are not four replications across independent donor samples. The larger estimate after excluding B indicates heterogeneous influence among preparations; it does not identify the reason for that heterogeneity or establish the pooled preparation’s identity.

All eight module-specific tests survived the eight-test correction family, with q-values from 0.001600 to 0.004300 (Figure 4A; Supplementary Data 1). The within-module incremental R^2^ ranged from 0.133295 for glycolysis/pentose-phosphate to 0.481138 for differentiation-associated proteins. Redox, proteostasis and the partial resolution panel had values of 0.453019, 0.416837 and 0.410559; mitochondrial, lipid and inflammasome/cytokine-related panels had values of 0.325271, 0.267469 and 0.230443. These proportions use different module-specific denominators. Their ordering is consequently not a ranking of contributions to the full multivariate effect, and the tests do not establish a common direction of change across conditions.

**Figure 4.**
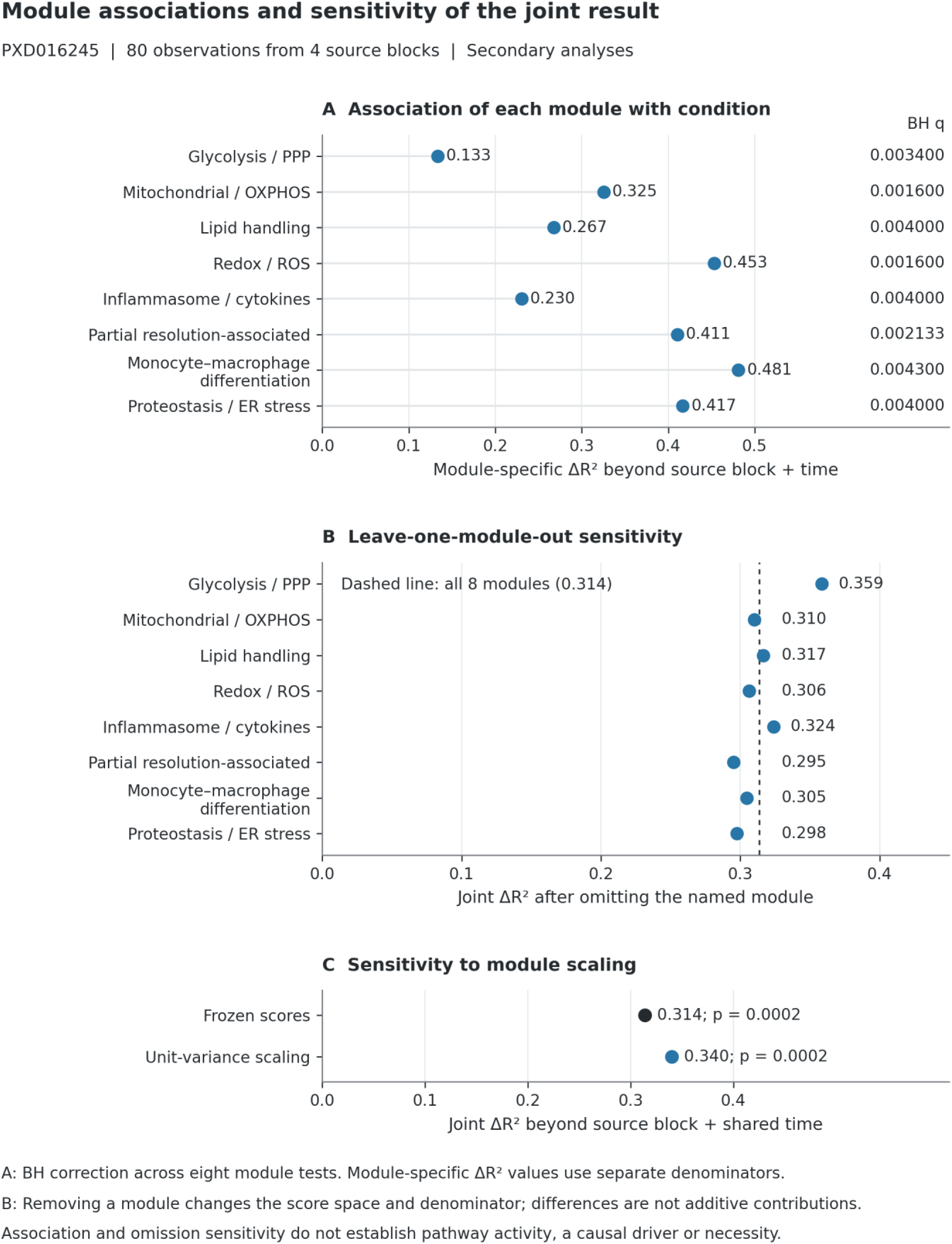
Module-specific associations and sensitivity of the joint result to score composition and scaling. All panels use PXD016245, with 80 repeated observations from four source blocks. (A) Frozen module-specific ΔR^2^ estimates for the joint condition-associated block beyond source block and shared time; q values are Benjamini– Hochberg-adjusted over eight module tests. Each ΔR^2^ uses the total variation of that module and is not a percentage contribution to the joint eight-module result. (B) The joint ΔR^2^ after omitting the named module, with the all-eight-module value (0.313663) shown as a dashed reference. Omission changes both the score space and its total-variation denominator; these differences are not additive contributions and do not identify necessary or causal drivers. (C) The joint result using the original frozen scores and after unit-variance scaling of the module columns. The scaling analysis is a secondary sensitivity analysis, not a replacement primary result. Association, omission sensitivity and scaling stability do not establish pathway activity or function. Points are frozen estimates without confidence intervals. Figure source data are provided in Supplementary Data 2.

Omitting any one module retained the multivariate association (all p = 0.0002; Figure 4B). The full-minus-omitted incremental R^2^ change ranged from −0.044889 for glycolysis/pentose-phosphate to 0.018480 for the partial resolution panel. The joint effect thus survived each single-module omission. An increase in incremental R^2^ after dropping glycolysis does not imply that glycolysis inhibits differentiation: removal changes the measured response space and its total variance. Equalizing module variances also retained the association (incremental R^2^ = 0.339754; pseudo-F = 4.403193; p = 0.0002). This is evidence of robustness to that scaling choice, while condition-dependent dispersion remains untested.

## 4 Discussion

### 4.1 What the adjusted association adds

The principal finding is that eight predefined protein-module scores retain a joint association with differentiation and polarization conditions after accounting for source block and shared differentiation time. Its magnitude, an additional 31.37% of the observed total score variation, and its persistence after each source block omission support a within-dataset description. The adjusted comparison adds to a visual account of the source differentiation proteome by specifying the covariates, response representation and exchangeability structure behind the association. It does not establish a new cell classification or an independently validated functional state.

The explicit separation of shared day effects from condition-associated terms is central to this interpretation. A broad increase in glycolysis-related scores was observed across conditions, whereas mitochondrial and proteostasis profiles differed in their terminal ordering. Such patterns can coexist in the same dataset. A single pooled contrast would describe their mixture without identifying the portion associated with condition after shared day variation is represented. Conversely, the joint block used here does not isolate a pure condition effect: its interaction component remains part of the primary estimate. The secondary interaction analysis provides supporting information but does not alter that definition.

### 4.2 Descriptive protein profiles differ across operational culture labels

The descriptive profiles are biologically informative because they identify specific mismatches between measured protein summaries and stereotyped functional expectations. M2-labelled cultures had the lowest terminal mitochondrial score, and the three-protein resolution-associated score was highest in M1-labelled cultures. These findings should be examined as properties of the culture system, measured features and scoring definitions. Reassigning cell identities on this basis would substitute one indirect annotation for another. Functional heterogeneity across macrophage stimuli is well established in primary experimental studies (1), but the present data do not demonstrate that a particular respiratory, inflammatory or efferocytic function followed the module scores.

The sign conventions make this caution concrete. A lower redox score reflects the relative abundances of positively and negatively oriented proteins within the panel. It cannot distinguish changes in oxidant production, antioxidant capacity, protein turnover or other mechanisms without appropriate measurements. Likewise, the resolution-associated score omits three intended members and shares APOE with lipid handling in the opposite orientation. Its elevation in M1-labelled cultures argues for describing the measured three-protein pattern directly. A functional explanation would require evidence beyond these abundance summaries.

The same distinction applies to the claim that all modules were associated with condition. The eight tests show breadth within the selected representation, but they do not demonstrate eight independently discovered pathways. Biological covariation may link panels, APOE is shared by two, and the protein sets were chosen before the revised test. No single omission abolished the association, yet leave-one-module-out analysis is not a perturbation experiment and cannot establish whether a biological process is necessary or dispensable. These are complementary statistical summaries of the same small dataset.

### 4.3 Temporal evidence supports a sampling window rather than an onset claim

Adjusted differences were detected from day 4 onward, whereas the day-2 test did not meet the correction threshold. This suggests that later sampling is informative for describing the studied protein configurations. It does not locate a transition between days 2 and 4, because an undetected early difference can reflect limited power or greater source block variability. The estimates also decreased between days 4 and 6, so the results do not support a steadily increasing separation. Testing a specific onset or trajectory shape would require an analysis designed for that endpoint and appropriately resolved early sampling.

Serial bulk culture measurements further limit what longitudinal language can imply. The observations represent samples linked by source block, condition and time; they are not repeated measurements of the same individual cells. Changes in the average proteome can therefore include shifts in cellular composition or survival as well as remodeling within cells. The block-preserving permutation scheme respects the available repeated structure, but cannot resolve those alternatives. The results are best understood as temporal profiles of source-matched cultures.

### 4.4 External validation remains unresolved

The ST001835 audit identified nonspecific chemical mappings and unresolved donor/time dependence. Its archived calculation therefore supplies neither positive nor negative biological evidence, and a curated, design-aware reanalysis would be required before interpreting it. The failure of this secondary comparison leaves the proteomic endpoint without external validation; its defects do not alter the verified PXD016245 calculation.

Independent macrophage metabolomic and lipidomic findings (4) (5) (6) remain biological context. They cannot supply validation merely because their subject matter is related. Testing portability would require a compatible independent dataset with the relevant culture contrasts and measurable module members, using a specified mapping and endpoint. Connecting protein abundance to function would additionally require suitable functional or metabolic measurements. Neither condition is satisfied by internal resampling or literature agreement alone.

### 4.5 Limitations and implications for reuse

The most consequential limitation is the mixture of three individual-donor preparations and one pooled preparation in four source blocks. Eighty serial observations cannot support population-wide claims about human myeloid differentiation. The pool is one composite source series, not 40 independent measurements, and its relationship to the individually obtained preparations is incompletely documented. Equal weighting of the four series is not an estimate across individual donors. Block-omission analyses reuse overlapping observations and baseline-inclusive standardized scores; they do not estimate transportability or remove all possible influence of pooled material on preprocessing. Confidence intervals for the multivariate effect were not generated, and descriptive ranges should not be mistaken for inferential intervals.

Technical and representational uncertainty also remain. The reanalysis reused source protein-group intensities and historical feature assignments. It did not independently validate peptide identifications, reprocess spectra, collapse multiple features per protein, or change the frozen scores. Partial protein coverage and the repeated contribution of G6PD and PDIA3 influence how the scores weight the measured proteome. The baseline-inclusive standardization defines their scale, so zero is not a biological reference state. Condition-linked acquisition or culture effects cannot be excluded by covariate adjustment alone. Restricted dispersion testing was not completed; therefore the multivariate association cannot be attributed exclusively to centroid differences independent of dispersion.

Prespecification and reproducibility help assess these choices but do not remove their limitations. The time-adjusted plan was fixed before its execution, after prior analyses of the same resource. The distinction is relevant when evaluating how much analytical independence the result has. Numerical reproduction confirms that the stored inputs and procedures yield the reported statistics; it does not establish biological validity or guarantee that the chosen representation is optimal. The revised figures and complete source tables make the observations, secondary-analysis withdrawal and sensitivity analyses available for that assessment.

Within these boundaries, PXD016245 supports a measurable association between culture condition and longitudinal proteomic module configurations beyond shared source block and day effects. The most informative extension is to test the specific observed protein patterns in an independent compatible system, while measuring the functions their annotations suggest. Until then, the result remains a descriptive account of source-matched human monocyte-derived cultures.

## Supporting information

Supplementary Data 3 - Reproducibility

## Data availability statement

The primary source data are publicly available in PRIDE/ProteomeXchange under accession PXD016245. The archived secondary dataset is available in Metabolomics Workbench under accession ST001835. The supplementary files provide the frozen score matrix, module definitions, results, descriptive figure data and an independent verifier of the primary endpoint. Verification starts from frozen scores; the source raw spectra remain available through the original proteomic accession.

## Ethics statement

This study used publicly accessible, de-identified secondary datasets. It involved no new participant recruitment, intervention, specimen collection or access to direct personal identifiers. The original PXD016245 sample collection was approved by the Sanquin Ethical Advisory Board, and the source investigators reported written informed consent from participants (3, p. 94). This attribution concerns the original collection; it is not a new approval number or institutional exemption for the present secondary analysis. Original approvals and consent procedures for other deposited source material remain attributable to the investigators and their source documentation.

## Author contributions

RN: Methodology, Validation, Visualization, Investigation, Writing – review and editing. KE: Validation, Investigation, Writing – review and editing. IKL: Validation, Writing – review and editing. AJ: Validation, Investigation, Writing – review and editing. YD-O: Investigation, Writing – review and editing. NG-D: Investigation, Supervision, Writing – review and editing. CF-P: Investigation, Supervision, Writing – review and editing. EZ: Investigation, Writing – review and editing. YB: Investigation, Writing – review and editing. LiP: Methodology, Resources, Validation, Investigation, Writing – review and editing. LeP: Methodology, Validation, Investigation, Writing – review and editing. AA: Methodology, Validation, Investigation, Supervision, Writing – review and editing. NG: Resources, Methodology, Validation, Investigation, Writing – review and editing. EN: Conceptualization, Methodology, Formal analysis, Data curation, Investigation, Validation, Visualization, Writing – original draft, Writing – review and editing, Supervision, Project administration, Funding acquisition.

## Acknowledgments

The authors acknowledge the investigators who generated and publicly deposited the source datasets, and Universidad Simón Bolívar, the Center for Research in Critical Dynamics and the Colombian Association of Immunology for their institutional and academic support.

## Generative artificial intelligence statement

Generative AI tools, including Claude Code and OpenAI Codex, assisted code review, literature organization, manuscript editing, figure scripting and numerical quality checks. OpenAI Codex based on GPT-6 assisted the September 2026 editorial revision and reconstruction of figures from existing data. No new experimental measurements or simulated biological observations were introduced. Scientific interpretation, authorship and responsibility for the submitted article remain with the authors.

