## Supplementary Methods for "Longitudinal proteomic module profiles vary across human monocyte differentiation and polarization conditions"

### Supplementary methods for longitudinal proteomic modules in human monocyte differentiation

#### Scope and analytical units

This supplement accompanies the reanalysis of PXD016245. Four filename-defined source blocks (A/B/C/P) each contribute serial bulk observations from four culture conditions at five post-baseline days, giving 80 observations and eight frozen scores. The original source describes three individual-donor preparations and one preparation pooled from 40 donors (Béguin, 2020, chapter 4, pp. 94–95). The four blocks must not be described as four individual donors. The inspected metadata do not explicitly establish which prefix denotes the pool. Legacy field names such as donor_id are retained in frozen files for traceability and denote source block. Supplementary Data 1 contains definitions and frozen statistics; Supplementary Data 2 contains descriptive summaries; Supplementary Data 3 verifies the original endpoint and documents this source-design correction.

#### Original protein identification and quantification

The public PXD016245 record identifies an Orbitrap Fusion instrument. The archived mqpar.xml file identifies MaxQuant version 1.5.3.30 and a search database named HUMAN170307.fasta. The exact database release and sequence count were not established from this parameter file. The original acquisition and identification workflow is distinguished from the secondary analysis: no raw spectra were reprocessed in the present work.

| Original search parameter | Deposited setting |
|---|---|
| Peptide and protein false-discovery thresholds | 0.01 and 0.01 |
| Enzyme and missed cleavages | Trypsin/P; maximum 2 |
| Fixed modification | Carbamidomethylation of cysteine |
| Variable modifications | Methionine oxidation; protein N-terminal acetylation |
| Initial and main precursor tolerance | 20 and 4.5 ppm |
| Match between runs | Enabled; separate MBR FDR option disabled |
| LFQ normalization | Enabled; minimum ratio count 2 |
| Minimum peptide length | 7 residues |
| Minimum peptides, razor peptides and unique peptides | 1, 1 and 0 |
| Contaminants and decoys in original search | Contaminants included; reversed decoys |

These parameters do not establish that each reported protein had two unique peptides. They are reproduced from the deposited search file and are not new filtering criteria selected for the revision. The primary experimental attribution is the monocyte-differentiation chapter in Béguin EP, Beneath the surface: Cellular responses in the vasculature, Utrecht University (2020), doi:10.33540/322, together with the dataset accession. A corresponding peer-reviewed source article was not independently verified. The unrelated DOI 10.1016/j.isci.2020.101074, present in some aggregated dataset metadata, is not used as the source citation.

#### Protein-group filtering and score construction

The source proteinGroups.txt contained 6,375 rows and 84 LFQ sample columns. Rows with a plus sign in Reverse, Potential contaminant or Only identified by site were excluded. The three flags occurred in 80, 144 and 149 rows, respectively, with overlap. Their union removed 346 rows and retained 6,029 protein-group features. The flag counts therefore must not be summed as mutually exclusive categories.

Only positive numeric LFQ values were log2 transformed. Other values were missing; no pseudocount or imputation was used. Each feature was standardized using observed values across the original 84 samples and the sample standard deviation (denominator n observed − 1; ddof = 1). Mapping and scoring then required at most 50% missing values across those samples and nonzero observed variance. The source LFQ normalization was inherited; no additional sample normalization or batch correction was applied at this stage.

All used protein memberships were obtained through exact normalized gene-symbol lookup. There were 54 distinct used assay features and 55 feature-module memberships. The only feature shared across two modules was APOE, with opposite signs in the lipid-handling and partial resolution panels. G6PD contributed two eligible features to glycolysis/pentose-phosphate and PDIA3 contributed two to proteostasis. A score was the arithmetic mean of the available sign-oriented feature z-scores, with at least two required. Multiple features for the same gene were not collapsed, so the implementation weights retained assay features rather than unique named proteins equally.

All eight scores were available for all 84 samples. The four day-0 monocyte observations were subsequently excluded from the condition analysis, leaving a complete 80 by 8 response matrix without rescoring. Consequently, the scores retain the baseline-inclusive standardization scale. A score of zero is a relative value on that scale, not a biological null or a zero-activity state.

#### Statistical definitions and uncertainty

The reduced model includes source block and categorical day. The full model adds condition and the day-by-condition interaction. Design ranks are 8 and 23, giving 15 degrees of freedom for the tested condition block and 57 residual degrees of freedom. SSblock equals SSEreduced minus SSEfull. The pseudo-F statistic is (SSblock/15)/(SSEfull/57), and incremental $R^2$ is SSblock/SStotal. These definitions test the joint condition block, not a condition main effect in isolation.

One condition-label assignment is applied to all five days within each source block. The 9,999 distinct non-identity assignments are sampled from the $24^4$ possible global assignments. The p-value includes a plus-one correction and a $10^{-9}$ statistic tolerance. The archived list is retained so that the result can be reproduced without drawing a new Monte Carlo sample. Independent reconstruction found one permuted statistic at least as large as the observed statistic, giving $p = 0.0002$. The verification reproduces the calculation from frozen scores; it does not replace validation of raw spectra, module biology or independent donor replication.

Leave-one-source-block-out fits use 60 observations and three source blocks, with design ranks 7 and 22, 15 degrees of freedom for the tested condition block and 38 residual degrees of freedom. Secondary day-specific analyses use 16 observations per day and separate total-variation denominators; their degrees of freedom for condition and residuals are 3 and 9. The five day-specific p-values and eight module-specific p-values form separate Benjamini–Hochberg families. The interaction sensitivity uses additive-model residual trajectories and is not described as a finite-sample exact test.

No bootstrap confidence intervals were calculated. Source-block ranges are descriptive minima and maxima among four heterogeneous preparations. The means weight each preparation equally, not each contributing person; the pool does not supply 40 separately measured donor outcomes. A range across leave-one-source-block-out effect estimates is not a confidence interval because the subsets overlap. Single-module omission changes both the multivariate response and its variance denominator. Neither module-specific incremental $R^2$ nor omission sensitivity identifies a causal driver.

Restricted dispersion testing was not implemented, so the primary association cannot be assigned exclusively to group-centroid differences independent of dispersion. The planned two-protein LRP1/TGFB1 sensitivity excluding APOE was also not implemented. These omissions are preserved rather than retrospectively substituted with new analyses. The historical blocked-sensitivity table is included in Supplementary Data 1.

The frozen P-exclusion sensitivity retained 60 observations and yielded pseudo-F = 2.946285, incremental $R^2$ = 0.315012 and permutation p = 0.002000. P is not explicitly equated with the pool. Moreover, the retained scores inherit standardization across all 84 original observations, so exclusion at the model stage is not a new preprocessing analysis free of pooled material. The possible overlap between the pool's contributors and individually obtained preparations was not established. Numerical reproducibility of the block-restricted permutations does not establish a donor-population sampling distribution.

#### Descriptive additions during revision

Condition-by-day summaries report n, mean, sample standard deviation, minimum and maximum for each module. There are 160 such cells, each based on four source blocks. The 32 module-condition summaries of day-10-minus-day-2 change preserve pairing within source block and report the number of positive and negative changes. These are descriptive additions made after the primary analysis was completed; no pairwise hypothesis tests, new module definitions or alternative inferential thresholds were introduced. Full summaries are supplied so that examples selected for the text are not the only accessible observations.

#### ST001835 historical analysis and withdrawal of interpretation

The source ST001835 study is Anders CB and colleagues, Journal of Leukocyte Biology (2022) 111:667–693, doi:10.1002/JLB.6A1120-744R. Its experimental findings should be distinguished from the archived secondary calculation described here. The historical secondary input comprised 498 features and 36 sample columns containing deposited area values. A maximum-missingness criterion of 50% retained 472 features and 35 samples. Sample M2b_028_070518 was removed because 252 of 498 entries were missing. The original paper's analysis used cell-count normalization, but that operation was not established for the deposited matrix reused by the secondary pipeline.

The retained matrix was log2 transformed and feature-wise standardized across 35 samples without imputation, additional sample normalization or batch correction. The historical scores used eight features annotated to glycolysis/pentose-phosphate, five to a mitochondrial/TCA panel and 49 to a lipid panel. The latter were all broad lipid-class matches. Unrestricted group-label permutations pooled observations annotated at 24 and 72 hours without donor or time adjustment. Donor parsing was recorded with medium confidence and included repeated identifiers.

Audit of the used feature ledger identified nonspecific assignments, including glucose and phosphate to glucose-6-phosphate, and substituted lactates to lactate. These assignments do not establish chemical identity. Together with unresolved label exchangeability, they preclude a biological interpretation of the historical p-values. The full used-feature audit and original numerical outputs are supplied for transparency. They are not evidence for or against macrophage metabolic differences and do not validate the PXD016245 endpoint.

Table S1 preserves the historical numerical calculations without endorsing their biological interpretation.

| Archived calculation | Statistic | Historical p-value | Interpretation after audit |
|---|---|---|---|
| Energy/DISCO-like | 5.198794 | 0.9268 | Not biologically interpretable under audited mapping and dependence assumptions |
| PERMANOVA | pseudo-F 0.679080; $R^2$ 0.104811 | 0.6826 | Not biologically interpretable under audited mapping and dependence assumptions |

No replacement metabolomic result is asserted in this revision. Reanalysis would require chemically curated identifiers, a verified donor/time design and an explicitly specified new analysis. The detected metabolite-name matching defects do not apply to the separately verified exact protein mapping in PXD016245.

#### Supplementary data contents

Supplementary Data 1 provides module definitions, protein and assay-feature memberships, missingness information, frozen primary and secondary proteomic statistics, unimplemented sensitivities and the documented historical ST001835 audit. Supplementary Data 2 provides the complete descriptive trajectory and paired-change summaries. Supplementary Data 3 supplies a self-contained independent verifier of the primary result from frozen scores and archived permutations, its required inputs, the historical plan and provenance records. Each archive includes a README specifying its scope. Reproduction from the score matrix should not be described as raw-spectra reanalysis or independent experimental replication.
