## Supplementary Data 1 for "Longitudinal proteomic module profiles vary across human monocyte differentiation and polarization conditions": READ_FIRST.md

Historical calculations retained for provenance only. Nonspecific metabolite-name mappings and unrestricted permutations despite donor/time structure preclude biological interpretation. Do not interpret the archived decision or p-values as validated negative evidence. See Supplementary Methods. No replacement test was performed.
