## Supplementary Data 1 for "Longitudinal proteomic module profiles vary across human monocyte differentiation and polarization conditions": README.md

Supplementary Data 1: frozen proteomic result tables and module/feature definitions. Statistics are historical, not newly selected results. Baseline input and permutations are in Supplementary Data 3. The explicitly labelled historical_ST001835_not_interpretable subfolder preserves withdrawn calculations solely for provenance; Supplementary Methods governs their interpretation.

SOURCE DESIGN CORRECTION (7 September 2026): the PXD016245 source study describes three individual-donor preparations and one preparation pooled from 40 donors. The filename prefixes A/B/C/P identify four source blocks, not four individual donors. Legacy fields such as donor_id and donor_excluded, LODO filenames, and donor wording in the archived tables retain their original bytes for traceability; in current interpretation they refer to source blocks. The mapping of P to the pooled preparation is inferred, not explicitly verified. The pool contributes one measured preparation, not 40 independently measured replicates. Within-block condition-trajectory permutations and all frozen statistics are unchanged. See SOURCE_DESIGN_CORRECTION.md and Supplementary Methods for the current interpretation.
