## Supplementary Data 1 for "Longitudinal proteomic module profiles vary across human monocyte differentiation and polarization conditions": SOURCE_DESIGN_CORRECTION.md

# Correction to biological-replication metadata

Correction date: 7 September 2026. This note updates interpretation and documentation; the archived scores, permutation assignments, numerical results, original analysis source and original preregistration are unchanged.

## What the available sources establish

The source study describes three individual-donor preparations plus one preparation pooled from 40 donors, measured during monocyte-derived differentiation. See Béguin (2020), chapter 4, printed pages 94–95, in the [Utrecht University thesis](https://research-portal.uu.nl/files/75476284/200915_thesis_eelke_beguin_upload_5f60b34aaace1.pdf). That chapter identifies the PXD016245 deposition. The source therefore does not support describing the experiment as four individual donors.

The original PXD016245 MaxQuant parameter file, `mqpar.xml`, lists the four filename prefixes A, B, C and P. Each occurs in the four condition series at days 2, 4, 6, 8 and 10, with one additional baseline monocyte sample. The parameter file does not explicitly define P as the pool or identify the individual contributors to any preparation.

The local historical importer, `src/cosmos_myeloid/prepare_pxd016245.py`, function `parse_sample_column` (lines 582–609 in the source inspected), captured a prefix matching `[ABCP]` from the sample-column name and assigned it to `donor_id`. The historical readiness report explicitly described these as donor-like prefixes and stated that no external sample sheet was used. Thus, the archived column is a filename-derived source-block identifier, not independently validated individual-donor metadata.

The correspondence **P = pooled preparation is a plausible inference from the labels and source design, not an explicitly verified sample-level mapping**. A, B and C must likewise not be described as individually verified donor identities solely on the basis of this parser. The source establishes the aggregate design of three individual-donor preparations and one pool; the available mapping evidence does not establish which block is which beyond inference.

## Consequences for the frozen analysis

- Refer to A/B/C/P as four **source blocks** or four **biological preparations**, not four independent individual donors. The primary matrix has 80 serial culture observations, with four observations per condition/day. A preparation pooled from 40 donors contributes one observed preparation; its contributors are not 40 independently measured analytical replicates.
- The original column name `donor_id` and historical code variable names remain intact for traceability. In the revised interpretation they denote source block. Fixed adjustment for that column is source-block adjustment.
- The archived permutation scheme exchanges the four condition trajectories **within each source block**, using the same exchange across all five days. It never exchanges observations between A, B, C and P. Relabeling the unit in the documentation does not change model matrices, ranks, the 24^4 assignment space, archived assignments, F statistics, incremental R2 or p values.
- Numerical reproducibility does not establish biological exchangeability, independence or representativeness. Interpretation remains conditional on the available preparations and the within-block exchangeability assumption. Pooling may change variability relative to individual-donor preparations, and overlap between pool contributors and the individual preparations is not resolved by the available metadata. Generalization to an individual-donor population is correspondingly limited.
- Refer to the historical leave-one-donor-out analysis as **leave-one-source-block-out** in current prose. If discussing P specifically, say exclusion of the P-labelled block, with its pooled status explicitly described as inferred.

## Already archived P-exclusion sensitivity

No new statistical analysis was performed for this correction. The existing table `reports/tables/PXD016245_time_aware_lodo.tsv` in the historical PXD016245 analysis contains the following row:

| Excluded source block | Observations | Reduced/full rank | Numerator/residual df | Pseudo-F | Incremental R2 | Permutation p |
| --- | ---: | --- | --- | ---: | ---: | ---: |
| P | 60 | 7 / 22 | 15 / 38 | 2.946285 | 0.315012 | 0.002000 |

This archived sensitivity used 9,999 sampled trajectory-preserving assignments within the three retained source blocks and met the historical joint criterion. Its source is outside the compact primary-verification inputs; this package reports the archived row for context and does **not** independently recompute that secondary result. The frozen primary scores were standardized on the original 84 samples, so row exclusion does not remove the P observations from the earlier feature-standardization step.

These results show that the archived condition-associated finding persisted after omission of the P-labelled block. They do not independently prove the sample-to-pool mapping, establish population-level replication, or constitute a newly performed donor-only analysis.

## Suggested interpretation

“The dataset comprised four source preparations: three individual-donor preparations and one preparation pooled from 40 donors. We retained the archived source-block labels A, B, C and P and adjusted for source block and time. Condition trajectories were permuted within each block while preserving day. The P-labelled block is presumed to represent the pool, although that mapping was not explicitly documented in the metadata available to us. In the previously archived sensitivity excluding P, the condition-associated result persisted (60 observations; pseudo-F = 2.946285; incremental R2 = 0.315012; permutation p = 0.002). The small number and mixed composition of preparations limit inference to individual-donor populations.”
