## Supplementary Data 2 for "Longitudinal proteomic module profiles vary across human monocyte differentiation and polarization conditions": README.md

Supplementary Data 2: descriptive summaries and figure source data derived from frozen scores. The four source blocks are identified as A, B, C and P. The original study describes three individual-donor preparations and one preparation pooled from 40 donors; the preparation-type-to-key mapping remains unverified. Means, sample standard deviations, observed ranges and paired day10-minus-day2 changes summarize four source blocks with equal weight, not four individual donors. The n column in the 160-cell summary counts source blocks. No pooled constituent is treated as a separately measured replicate. Figure 4 omission data report the joint delta_R2 WITHOUT the named module, not an additive contribution. Legacy donor wording in certain filenames is retained for link compatibility only; revised column labels and this description define their interpretation.
