## Supplementary Data 3 - Reproducibility for "Longitudinal proteomic module profiles vary across human monocyte differentiation and polarization conditions": pxd016245_time_aware_audit.py

```
"""Independent audit of the executed A001-R5 time-aware analysis (A004).

This module deliberately does NOT import the A003 execution functions. It re-derives Y
and the design matrices with a DIFFERENT reference coding (drop-last instead of
drop-first) so that agreement proves the inference depends on the design column space, not
on an arbitrary coding. The primary p-value is recomputed from the FROZEN permutation
registry written by A003 (not a freshly generated list). LODO/day/module/interaction are
reproduced via the same documented deterministic generator and compared to the A003 tables.

Classification: AUDIT_PASS / AUDIT_PASS_WITH_DOCUMENTATION_CORRECTION / AUDIT_FAIL.
"""
from __future__ import annotations

import csv
import hashlib
import itertools
import os

import numpy as np

MODULES = [
    "glycolysis_ppp", "mitochondrial_oxphos", "lipid_foam_cell", "redox_ros",
    "inflammasome_cytokine", "resolution_efferocytosis",
    "monocyte_to_macrophage_differentiation", "proteostasis_er_stress",
]
DONORS = ["A", "B", "C", "P"]
TIMEPOINTS = [2, 4, 6, 8, 10]
PHENO = ["M1", "M2", "dendritic_cell", "osteoclast"]
PERMS = list(itertools.permutations(range(4)))
SEED = 20260615
B = 9999
TOL = 1e-9
NUM_TOL = 1e-9
STORED_TOL = 1e-6   # tables store 6-decimal-rounded values; compare to that precision
INPUT_SHA = "1fb757edd9bbb8e2354547ab733fc04a7888cad661b19e488a6e13c4664a8e66"
PREREG_SHA = "ce2d30280a7619b1d7c7f74913d6d616e90819682f7025cfe3949038a0d15ef6"


def _sha(path):
    """Raw working-copy SHA-256 (used to verify the SEALED references 1fb757/ce2d30,
    which were sealed on this autocrlf=true Windows working copy)."""
    return hashlib.sha256(open(path, "rb").read()).hexdigest()


def _sha_lf(path):
    """LF-normalized SHA-256 == the git blob hash. Portable and reproducible on any
    platform regardless of autocrlf; used for the A004 freeze manifest/artifact hashes."""
    return hashlib.sha256(open(path, "rb").read().replace(b"\r\n", b"\n")).hexdigest()


def _read_input(path):
    rows = list(csv.DictReader(open(path, encoding="utf-8"), delimiter="\t"))
    donor = np.array([r["donor_id"] for r in rows])
    pheno = np.array([r["phenotype"] for r in rows])
    tp = np.array([int(r["timepoint"]) for r in rows])
    Y = np.array([[float(r[m]) for m in MODULES] for r in rows], dtype=float)
    return donor, pheno, tp, Y


# --- INDEPENDENT design: drop-LAST reference coding (A003 used drop-first) ---
def _dummies_drop_last(values, categories):
    cols = [(values == c).astype(float) for c in categories[:-1]]
    return np.column_stack(cols) if cols else np.empty((len(values), 0))


def _codes(pheno):
    idx = {p: i for i, p in enumerate(PHENO)}
    return np.array([idx[p] for p in pheno])


def _pheno_oh(codes):  # drop-last: columns for codes 0,1,2 (ref = osteoclast=3)
    return np.column_stack([(codes == k).astype(float) for k in (0, 1, 2)])


def _fixed(donor, tp):
    n = len(donor)
    return np.hstack([np.ones((n, 1)), _dummies_drop_last(donor, DONORS),
                      _dummies_drop_last(tp, TIMEPOINTS)])


def _time_d(tp):
    return _dummies_drop_last(tp, TIMEPOINTS)


def _full(fixed, time_d, codes):
    ph = _pheno_oh(codes)
    inter = (time_d[:, :, None] * ph[:, None, :]).reshape(len(codes), -1)
    return np.hstack([fixed, ph, inter])


def _additive(fixed, codes):
    return np.hstack([fixed, _pheno_oh(codes)])


def _sse(Z, Y):
    beta = np.linalg.lstsq(Z, Y, rcond=None)[0]
    r = Y - Z @ beta
    return r, float(np.sum(r ** 2))


def _f(ssb, ssef, dfb, dfr):
    return (ssb / dfb) / (ssef / dfr)


def _gen_assign(n_donors, b):
    rng = np.random.Generator(np.random.PCG64(SEED))
    total, identity, seen, out = 24 ** n_donors, tuple([0] * n_donors), set(), []
    while len(out) < b:
        x = int(rng.integers(0, total))
        t, y = [], x
        for _ in range(n_donors):
            t.append(y % 24)
            y //= 24
        t = tuple(t)
        if t == identity or t in seen:
            continue
        seen.add(t)
        out.append(t)
    return out


def _perm_codes(codes, donor, donors_order, assign):
    out = codes.copy()
    for di, d in enumerate(donors_order):
        P = np.array(PERMS[assign[di]])
        m = donor == d
        out[m] = P[codes[m]]
    return out


def _bh(p):
    p = np.asarray(p, float)
    m = len(p)
    o = np.argsort(p)
    ranked = p[o] * m / np.arange(1, m + 1)
    q = np.minimum.accumulate(ranked[::-1])[::-1]
    out = np.empty(m)
    out[o] = np.clip(q, 0, 1)
    return out


def _table(tbl, name):
    with open(os.path.join(tbl, name), encoding="utf-8") as f:
        return list(csv.DictReader(f, delimiter="\t"))


def _kv(tbl, name):
    rows = _table(tbl, name)
    return {r[list(r)[0]]: r[list(r)[1]] for r in rows}


def audit(root, write=True):
    tbl = os.path.join(root, "reports", "tables")
    inp = os.path.join(tbl, "PXD016245_time_aware_analysis_input.tsv")
    prereg = os.path.join(root, "PXD016245_STATE_SEPARATION_TIME_AWARE_PREREG.md")

    cmp_rows = []   # (check, reported, recomputed, abs_diff, status)
    findings = []

    def add(check, reported, recomputed, ok, absdiff=""):
        cmp_rows.append([check, reported, recomputed, absdiff, "PASS" if ok else "FAIL"])
        if not ok:
            findings.append(check)

    # hashes
    add("input_sha256", INPUT_SHA, _sha(inp), _sha(inp) == INPUT_SHA)
    add("sealed_prereg_sha256", PREREG_SHA, _sha(prereg), _sha(prereg) == PREREG_SHA)

    donor, pheno, tp, Y = _read_input(inp)
    codes = _codes(pheno)
    n = len(donor)
    add("n_rows", "80", str(n), n == 80)
    add("factorial_complete", "True",
        str(len({(donor[i], pheno[i], tp[i]) for i in range(n)}) == 80
            and set(donor) == set(DONORS) and set(pheno) == set(PHENO)
            and set(tp.tolist()) == set(TIMEPOINTS)),
        len({(donor[i], pheno[i], tp[i]) for i in range(n)}) == 80)

    fixed, td = _fixed(donor, tp), _time_d(tp)
    ZR, ZF, ZA = fixed, _full(fixed, td, codes), _additive(fixed, codes)
    rR, rF, rA = (int(np.linalg.matrix_rank(ZR)), int(np.linalg.matrix_rank(ZF)),
                  int(np.linalg.matrix_rank(ZA)))
    add("rank_reduced", "8", str(rR), rR == 8)
    add("rank_additive", "11", str(rA), rA == 11)
    add("rank_full", "23", str(rF), rF == 23)

    SST = float(np.sum((Y - Y.mean(0)) ** 2))
    rRr, sseR = _sse(ZR, Y)
    rFr, sseF = _sse(ZF, Y)
    _, sseA = _sse(ZA, Y)
    SSb = sseR - sseF
    F_obs = _f(SSb, sseF, 15, 57)
    dR2 = SSb / SST
    R2r, R2f = 1 - sseR / SST, 1 - sseF / SST

    pr = _kv(tbl, "PXD016245_time_aware_primary_result.tsv")
    F_rep, dR2_rep = float(pr["F_observed"]), float(pr["delta_R2"])
    add("F_observed", f"{F_rep:.6f}", f"{F_obs:.6f}", abs(F_obs - F_rep) < STORED_TOL, f"{abs(F_obs-F_rep):.2e}")
    add("delta_R2", f"{dR2_rep:.6f}", f"{dR2:.6f}", abs(dR2 - dR2_rep) < STORED_TOL, f"{abs(dR2-dR2_rep):.2e}")
    add("delta_R2==R2f-R2r", f"{dR2:.6f}", f"{R2f-R2r:.6f}", abs(dR2 - (R2f - R2r)) < 1e-12)
    add("delta_R2==SSb/SST", f"{dR2:.6f}", f"{SSb/SST:.6f}", abs(dR2 - SSb / SST) < 1e-12)

    # independence: SSE under drop-last coding must equal A003's reported SSE_full (drop-first)
    add("SSE_full_coding_invariant", pr["SSE_full"], f"{sseF:.6f}",
        abs(sseF - float(pr["SSE_full"])) < 1e-6, f"{abs(sseF-float(pr['SSE_full'])):.2e}")

    # ---- primary p from the FROZEN registry ----
    reg = _table(tbl, "PXD016245_time_aware_permutation_registry.tsv")
    add("registry_rows", "9999", str(len(reg)), len(reg) == 9999)
    tuples = [(int(r["assign_A"]), int(r["assign_B"]), int(r["assign_C"]), int(r["assign_P"])) for r in reg]
    add("registry_unique", "9999", str(len(set(tuples))), len(set(tuples)) == 9999)
    add("registry_no_identity", "absent", "present" if (0, 0, 0, 0) in set(tuples) else "absent",
        (0, 0, 0, 0) not in set(tuples))
    add("registry_valid_indices", "0..23", "ok" if all(0 <= v < 24 for t in tuples for v in t) else "bad",
        all(0 <= v < 24 for t in tuples for v in t))

    maxFdiff = 0.0
    F_perm_audit = np.empty(len(reg))
    for k, (t, r) in enumerate(zip(tuples, reg)):
        cp = _perm_codes(codes, donor, DONORS, t)
        _, ssef = _sse(_full(fixed, td, cp), Y)
        Fp = _f(sseR - ssef, ssef, 15, 57)
        F_perm_audit[k] = Fp
        maxFdiff = max(maxFdiff, abs(Fp - float(r["F_perm"])))
    add("registry_F_perm_match", "match", f"max|d|={maxFdiff:.2e}", maxFdiff < 1e-6, f"{maxFdiff:.2e}")
    b = int(np.sum(F_perm_audit >= F_obs - TOL))
    p_audit = (b + 1) / (B + 1)
    p_rep = float(pr["p_primary"])
    add("primary_b_count", "", str(b), True)
    add("primary_p_value", f"{p_rep:.6f}", f"{p_audit:.6f}", abs(p_audit - p_rep) < NUM_TOL, f"{abs(p_audit-p_rep):.2e}")

    primary_supported = (p_audit <= 0.05) and (dR2 > 0.02)
    add("PRIMARY_SUPPORTED", pr["PRIMARY_SUPPORTED"], str(primary_supported),
        str(primary_supported) == pr["PRIMARY_SUPPORTED"])

    # ---- LODO (regenerate via documented generator) ----
    lodo_tbl = {r["donor_excluded"]: r for r in _table(tbl, "PXD016245_time_aware_lodo.tsv")}
    lodo_keep = True
    for d_excl in DONORS:
        keep = donor != d_excl
        dd, pp, tt, YY = donor[keep], pheno[keep], tp[keep], Y[keep]
        cc = _codes(pp)
        remaining = [d for d in DONORS if d != d_excl]
        fx = np.hstack([np.ones((len(dd), 1)), _dummies_drop_last(dd, remaining),
                        _dummies_drop_last(tt, TIMEPOINTS)])
        tdl = _dummies_drop_last(tt, TIMEPOINTS)
        rk_R = int(np.linalg.matrix_rank(fx))
        ZFl = _full(fx, tdl, cc)
        rk_F = int(np.linalg.matrix_rank(ZFl))
        _, sR = _sse(fx, YY)
        _, sF = _sse(ZFl, YY)
        SSTk = float(np.sum((YY - YY.mean(0)) ** 2))
        Fo = _f(sR - sF, sF, 15, 38)
        dR2o = (sR - sF) / SSTk
        a3 = _gen_assign(3, B)
        Fp = np.empty(B)
        for j, a in enumerate(a3):
            cp = _perm_codes(cc, dd, remaining, a)
            _, sf = _sse(_full(fx, tdl, cp), YY)
            Fp[j] = _f(sR - sf, sf, 15, 38)
        pj = (int(np.sum(Fp >= Fo - TOL)) + 1) / (B + 1)
        joint = (pj <= 0.05) and (dR2o > 0.02)
        lodo_keep = lodo_keep and joint
        rep = lodo_tbl[d_excl]
        ok = (rk_R == 7 and rk_F == 22 and abs(Fo - float(rep["F"])) < 1e-6
              and abs(pj - float(rep["p"])) < NUM_TOL and abs(dR2o - float(rep["delta_R2"])) < 1e-6)
        add(f"LODO_{d_excl}", f"F={rep['F']},p={rep['p']},dR2={rep['delta_R2']}",
            f"F={Fo:.6f},p={pj:.6f},dR2={dR2o:.6f},ranks={rk_R}/{rk_F}", ok)

    # ---- interaction (Freedman-Lane) ----
    rA_resid, _ = _sse(ZA, Y)
    Yhat_A = Y - rA_resid
    pos = {(donor[i], codes[i], tp[i]): i for i in range(n)}
    F_int_obs = _f(sseA - sseF, sseF, 12, 57)
    a4 = _gen_assign(4, B)
    Fip = np.empty(B)
    for j, a in enumerate(a4):
        src = np.arange(n)
        for di, d in enumerate(DONORS):
            P = PERMS[a[di]]
            for i in np.where(donor == d)[0]:
                src[i] = pos[(d, P[codes[i]], tp[i])]
        Ystar = Yhat_A + rA_resid[src]
        _, ssea = _sse(ZA, Ystar)
        _, ssef = _sse(ZF, Ystar)
        Fip[j] = _f(ssea - ssef, ssef, 12, 57)
    p_int = (int(np.sum(Fip >= F_int_obs - TOL)) + 1) / (B + 1)
    it = _table(tbl, "PXD016245_time_aware_interaction_sensitivity.tsv")[0]
    add("interaction_F", it["F_observed"], f"{F_int_obs:.6f}", abs(F_int_obs - float(it["F_observed"])) < 1e-6)
    add("interaction_p", it["p"], f"{p_int:.6f}", abs(p_int - float(it["p"])) < NUM_TOL)
    add("interaction_non_decisional", "yes",
        "yes" if "non_decisional" in it["role"] or "non-decisional" in it["role"].lower() else "no",
        ("non_decisional" in it["role"] or "non-decisional" in it["role"].lower()))

    # ---- day-specific BH ----
    day_tbl = {int(r["timepoint"]): r for r in _table(tbl, "PXD016245_time_aware_day_specific.tsv")}
    day_p = []
    for t in TIMEPOINTS:
        sel = tp == t
        dd, pp, YY = donor[sel], pheno[sel], Y[sel]
        cc = _codes(pp)
        fx = np.hstack([np.ones((len(dd), 1)), _dummies_drop_last(dd, DONORS)])
        ZFd = np.hstack([fx, _pheno_oh(cc)])
        dfb = int(np.linalg.matrix_rank(ZFd)) - int(np.linalg.matrix_rank(fx))
        dfr = len(dd) - int(np.linalg.matrix_rank(ZFd))
        _, sR = _sse(fx, YY)
        _, sF = _sse(ZFd, YY)
        Fo = _f(sR - sF, sF, dfb, dfr)
        a = _gen_assign(4, B)
        Fp = np.empty(B)
        for j, av in enumerate(a):
            cp = _perm_codes(cc, dd, DONORS, av)
            _, sf = _sse(np.hstack([fx, _pheno_oh(cp)]), YY)
            Fp[j] = _f(sR - sf, sf, dfb, dfr)
        day_p.append((int(np.sum(Fp >= Fo - TOL)) + 1) / (B + 1))
    day_q = _bh(day_p)
    maxq = max(abs(q - float(day_tbl[t]["q_bh"])) for t, q in zip(TIMEPOINTS, day_q))
    add("day_BH_q", "match", f"max|d|={maxq:.2e}", maxq < 1e-6, f"{maxq:.2e}")

    # ---- module-specific BH (reuse primary frozen registry codes) ----
    sseR_col = np.sum(rRr ** 2, 0)
    sseF_col = np.sum(rFr ** 2, 0)
    SST_col = np.sum((Y - Y.mean(0)) ** 2, 0)
    Fmod_obs = np.array([_f(sseR_col[m] - sseF_col[m], sseF_col[m], 15, 57) for m in range(8)])
    Fmod_perm = np.empty((8, len(reg)))
    for k, t in enumerate(tuples):
        cp = _perm_codes(codes, donor, DONORS, t)
        rf, _ = _sse(_full(fixed, td, cp), Y)
        sc = np.sum(rf ** 2, 0)
        Fmod_perm[:, k] = [_f(sseR_col[m] - sc[m], sc[m], 15, 57) for m in range(8)]
    pmod = np.array([(int(np.sum(Fmod_perm[m] >= Fmod_obs[m] - TOL)) + 1) / (B + 1) for m in range(8)])
    qmod = _bh(pmod)
    mod_tbl = {r["module"]: r for r in _table(tbl, "PXD016245_time_aware_module_specific.tsv")}
    maxmq = max(abs(qmod[m] - float(mod_tbl[MODULES[m]]["q_bh"])) for m in range(8))
    add("module_BH_q", "match", f"max|d|={maxmq:.2e}", maxmq < 1e-6, f"{maxmq:.2e}")

    # ---- leave-one-module-out sign documentation ----
    lomo = _table(tbl, "PXD016245_time_aware_leave_one_module_out.tsv")
    sign_ok = True
    for row in lomo:
        full = float(row["delta_R2_full_space"])
        without = float(row["delta_R2_without_module"])
        change = float(row["change_in_delta_R2"])
        if abs((full - without) - change) > 1e-9:
            sign_ok = False
    add("LOMO_sign_is_full_minus_without", "delta_R2_full - delta_R2_without",
        "confirmed" if sign_ok else "MISMATCH", sign_ok)

    # ---- scaling ----
    sd = Y.std(0, ddof=1)
    Ys = (Y - Y.mean(0)) / sd
    SSTs = float(np.sum((Ys - Ys.mean(0)) ** 2))
    _, sRs = _sse(ZR, Ys)
    _, sFs = _sse(ZF, Ys)
    Fscal = _f(sRs - sFs, sFs, 15, 57)
    sc = _table(tbl, "PXD016245_time_aware_scaling_sensitivity.tsv")[0]
    add("scaling_F", sc["F"], f"{Fscal:.6f}", abs(Fscal - float(sc["F"])) < 1e-6)
    add("scaling_non_decisional", "yes", sc["role"], sc["role"] == "supportive_non_decisional")

    # ---- blocked sensitivities still blocked ----
    bl = _table(tbl, "PXD016245_time_aware_blocked_sensitivities.tsv")
    names = {b_["sensitivity"] for b_ in bl}
    add("PERMDISP_blocked", "BLOCKED",
        "BLOCKED" if any("PERMDISP" in nme for nme in names) else "missing",
        any("PERMDISP" in nme for nme in names))
    add("LRP1_TGFB1_blocked", "BLOCKED",
        "BLOCKED" if any("APOE" in nme for nme in names) else "missing",
        any("APOE" in nme for nme in names))

    # ---- decision reproduced independently of sensitivities ----
    def decide(prim_ok, lodo_all_keep, design_ok=True,
               interaction=None, day10=None, scaling=None):
        # sensitivities are intentionally ignored
        if not design_ok:
            return "EXECUTION_BLOCKED_INVALID_DESIGN"
        if not prim_ok:
            return "ORIGINAL_SEPARATION_NOT_SUPPORTED_AFTER_TIME_ADJUSTMENT"
        if not lodo_all_keep:
            return "AMBIGUOUS_LOW_DONOR_STABILITY_C1"
        return "TIME_ADJUSTED_PHENOTYPE_ARCHITECTURE_SUPPORTED_C1"

    decision = decide(primary_supported, lodo_keep)
    # robustness: flipping any sensitivity must not change the decision
    perturbed = {decide(primary_supported, lodo_keep, interaction=x, day10=y, scaling=z)
                 for x in (True, False) for y in (True, False) for z in (True, False)}
    add("decision_ignores_sensitivities", "1 outcome", str(len(perturbed)), len(perturbed) == 1)
    fd = _kv(tbl, "PXD016245_time_aware_final_decision.tsv")
    add("final_decision_reproduced", fd["decision"], decision, decision == fd["decision"])

    # claim audit: no prohibited claims permitted
    ct = {r["claim"]: r["permitted"] for r in _table(tbl, "PXD016245_time_aware_claim_trace.tsv")}
    prohibited_ok = all(v == "no" for k, v in ct.items()
                        if any(w in k for w in ("mechanism", "flux", "validation", "therapeutic", "C2 / C3")))
    add("no_prohibited_claims_permitted", "all no", "ok" if prohibited_ok else "leak", prohibited_ok)

    classification = "AUDIT_PASS" if not findings else "AUDIT_FAIL"

    if write:
        _write_audit(tbl, root, cmp_rows, classification, dict(
        F_obs=F_obs, dR2=dR2, p=p_audit, b=b, decision=decision,
        F_int=F_int_obs, p_int=p_int, Fscal=Fscal), reg, inp, prereg)
    return dict(classification=classification, findings=findings, cmp_rows=cmp_rows,
                F_obs=F_obs, dR2=dR2, p=p_audit, b=b, decision=decision,
                day_q=[float(x) for x in day_q], qmod=[float(x) for x in qmod],
                p_int=p_int, F_int=F_int_obs, Fscal=Fscal)


def _w(path, header, rows):
    with open(path, "w", encoding="utf-8") as f:
        f.write("\t".join(header) + "\n")
        for r in rows:
            f.write("\t".join(str(x) for x in r) + "\n")


FREEZE_ARTIFACTS = [
    ("reports/tables/PXD016245_time_aware_analysis_input.tsv", "sealed locked input"),
    ("PXD016245_STATE_SEPARATION_TIME_AWARE_PREREG.md", "sealed preregistration"),
    ("reports/tables/PXD016245_time_aware_permutation_registry.tsv", "frozen 9999 permutations"),
    ("reports/tables/PXD016245_time_aware_primary_result.tsv", "primary result"),
    ("reports/tables/PXD016245_time_aware_lodo.tsv", "leave-one-donor-out"),
    ("reports/tables/PXD016245_time_aware_interaction_sensitivity.tsv", "interaction (non-decisional)"),
    ("reports/tables/PXD016245_time_aware_day_specific.tsv", "day-specific BH"),
    ("reports/tables/PXD016245_time_aware_module_specific.tsv", "module-specific BH"),
    ("reports/tables/PXD016245_time_aware_leave_one_module_out.tsv", "leave-one-module-out"),
    ("reports/tables/PXD016245_time_aware_scaling_sensitivity.tsv", "scaling (non-decisional)"),
    ("reports/tables/PXD016245_time_aware_blocked_sensitivities.tsv", "blocked sensitivities"),
    ("reports/tables/PXD016245_time_aware_final_decision.tsv", "final decision"),
    ("PXD016245_TIME_AWARE_SAFE_INTERPRETATION.md", "safe interpretation"),
    ("src/cosmos_myeloid/pxd016245_time_aware_execution.py", "execution module"),
    ("scripts/79_execute_pxd016245_time_aware.py", "execution script"),
    ("tests/test_pxd016245_time_aware_execution.py", "execution tests"),
]


def _write_audit(tbl, root, cmp_rows, classification, vals, reg, inp, prereg):
    _w(os.path.join(tbl, "PXD016245_time_aware_audit_comparison.tsv"),
       ["check", "reported", "recomputed", "abs_diff", "status"], cmp_rows)

    _w(os.path.join(tbl, "PXD016245_time_aware_decision_reproduction.tsv"),
       ["field", "value"],
       [["F_observed", f"{vals['F_obs']:.6f}"], ["delta_R2", f"{vals['dR2']:.6f}"],
        ["b_extreme_count", vals["b"]], ["p_primary", f"{vals['p']:.6f}"],
        ["decision", vals["decision"]],
        ["decision_inputs", "stop_rule + primary_criterion + four LODO only"],
        ["sensitivities_used_in_decision", "none"]])

    _w(os.path.join(tbl, "PXD016245_time_aware_frozen_values.tsv"),
       ["quantity", "frozen_value"],
       [["F_observed", f"{vals['F_obs']:.6f}"], ["delta_R2", f"{vals['dR2']:.6f}"],
        ["p_primary", f"{vals['p']:.6f}"], ["b", vals["b"]], ["B", B], ["seed", SEED],
        ["interaction_F", f"{vals['F_int']:.6f}"], ["interaction_p", f"{vals['p_int']:.6f}"],
        ["scaling_F", f"{vals['Fscal']:.6f}"], ["decision", vals["decision"]],
        ["input_sha256", INPUT_SHA], ["sealed_prereg_sha256", PREREG_SHA]])

    rows = []
    for rel, role in FREEZE_ARTIFACTS:
        path = os.path.join(root, rel)
        if not os.path.exists(path):
            rows.append([rel, "MISSING", "", "", role])
            continue
        size = len(open(path, "rb").read().replace(b"\r\n", b"\n"))
        rows.append([rel, _sha_lf(path), size, role, "true"])
    _w(os.path.join(tbl, "PXD016245_time_aware_artifact_hashes.tsv"),
       ["path", "sha256_lf_normalized", "size_bytes_lf", "role", "frozen"], rows)

    man = []
    for rel, role in FREEZE_ARTIFACTS:
        path = os.path.join(root, rel)
        if not os.path.exists(path):
            continue
        nr = nc = "NA"
        if rel.endswith(".tsv"):
            lines = [ln for ln in open(path, encoding="utf-8").read().splitlines() if ln]
            nr, nc = str(len(lines) - 1), str(len(lines[0].split("\t")))
        size_lf = len(open(path, "rb").read().replace(b"\r\n", b"\n"))
        man.append([os.path.basename(rel), rel, _sha_lf(path), size_lf,
                    nr, nc, role, "ff67ee8", "true",
                    "LF-normalized (git-blob) hash; audited and frozen by A004"])
    _w(os.path.join(tbl, "PXD016245_time_aware_result_manifest.tsv"),
       ["artifact", "path", "sha256_lf_normalized", "size_bytes_lf", "rows", "columns",
        "role", "source_commit", "frozen", "notes"], man)
```
