## Supplementary Data 3 - Reproducibility for "Longitudinal proteomic module profiles vary across human monocyte differentiation and polarization conditions": pxd016245_time_aware_execution.py

```
"""Execute the sealed A001-R5 PXD016245 time-aware analysis.

Confirmatory, deterministic implementation of the SEALED preregistration. It does NOT
re-score, re-standardize or modify the locked input; it reads the frozen 80x8 module
matrix and runs exactly the preregistered models, permutation scheme and decision tree.

Statistic (per prereg 7a): multivariate linear model on the 8-module matrix, equivalent
to Euclidean PERMANOVA. For a model with design Z (full-rank, intercept):
    beta  = numpy.linalg.lstsq(Z, Y, rcond=None)[0]
    SSE   = sum((Y - Z@beta)**2)            # squared Frobenius norm
    SST   = sum((Y - column_mean(Y))**2)
    SS_block = SSE_reduced - SSE_full
    F_block  = (SS_block / df_block) / (SSE_full / df_resid)
    delta_R2 = SS_block / SST
Permutation: trajectory-preserving within donor; canonical order M1, M2, dendritic_cell,
osteoclast; 24^4 space; identity excluded; 9999 distinct draws without replacement;
PCG64(20260615); tie rule F_perm >= F_observed - 1e-9; p = (b+1)/(B+1).
"""
from __future__ import annotations

import csv
import itertools
import os
from datetime import datetime, timezone

import numpy as np

MODULES = [
    "glycolysis_ppp", "mitochondrial_oxphos", "lipid_foam_cell", "redox_ros",
    "inflammasome_cytokine", "resolution_efferocytosis",
    "monocyte_to_macrophage_differentiation", "proteostasis_er_stress",
]
DONORS = ["A", "B", "C", "P"]
TIMEPOINTS = [2, 4, 6, 8, 10]
PHENO = ["M1", "M2", "dendritic_cell", "osteoclast"]   # canonical order (prereg 8a)
PERMS = list(itertools.permutations(range(4)))          # 24; PERMS[0] = identity
SEED = 20260615
B = 9999
TOL = 1e-9
INPUT_SHA256 = "1fb757edd9bbb8e2354547ab733fc04a7888cad661b19e488a6e13c4664a8e66"


# ---------- IO ----------

def load_input(path):
    rows = list(csv.DictReader(open(path, encoding="utf-8"), delimiter="\t"))
    donor = np.array([r["donor_id"] for r in rows])
    pheno = np.array([r["phenotype"] for r in rows])
    tp = np.array([int(r["timepoint"]) for r in rows])
    Y = np.array([[float(r[m]) for m in MODULES] for r in rows], dtype=float)
    return donor, pheno, tp, Y, rows


# ---------- design ----------

def _dummies(values, categories):
    """full-rank drop-first dummies (float), columns for categories[1:]."""
    cols = [(values == c).astype(float) for c in categories[1:]]
    return np.column_stack(cols) if cols else np.empty((len(values), 0))


def fixed_blocks(donor, tp):
    n = len(donor)
    intercept = np.ones((n, 1))
    donor_d = _dummies(donor, DONORS)
    time_d = _dummies(tp, TIMEPOINTS)
    fixed = np.hstack([intercept, donor_d, time_d])   # 8 cols on full data
    return fixed, time_d


def pheno_onehot(codes):
    return np.column_stack([(codes == k).astype(float) for k in (1, 2, 3)])


def build_full(fixed, time_d, codes):
    ph = pheno_onehot(codes)                                  # n x 3
    inter = (time_d[:, :, None] * ph[:, None, :]).reshape(len(codes), -1)  # n x (t*p)
    return np.hstack([fixed, ph, inter])


def build_additive(fixed, codes):
    return np.hstack([fixed, pheno_onehot(codes)])


def codes_of(pheno):
    idx = {p: i for i, p in enumerate(PHENO)}
    return np.array([idx[p] for p in pheno])


def sse(Z, Y):
    beta = np.linalg.lstsq(Z, Y, rcond=None)[0]
    resid = Y - Z @ beta
    return resid, float(np.sum(resid ** 2))


def colsumsq(resid):
    return np.sum(resid ** 2, axis=0)


# ---------- permutation sampling ----------

def sample_global(rng, n_donors, b):
    total = 24 ** n_donors
    identity = tuple([0] * n_donors)
    seen, out = set(), []
    while len(out) < b:
        x = int(rng.integers(0, total))
        t, y = [], x
        for _ in range(n_donors):
            t.append(y % 24)
            y //= 24
        t = tuple(t)
        if t == identity or t in seen:
            continue
        seen.add(t)
        out.append(t)
    return out


def permuted_codes(codes_obs, donor, donors_order, assign):
    """Relabel phenotype within each donor by its permutation (trajectory-preserving)."""
    out = codes_obs.copy()
    for di, d in enumerate(donors_order):
        P = np.array(PERMS[assign[di]])
        mask = donor == d
        out[mask] = P[codes_obs[mask]]
    return out


def residual_src(donor, pheno, tp, donors_order, assign):
    """Row index map for trajectory-preserving permutation of residuals (Freedman-Lane).

    target row (d, code c, t) <- source row (d, code PERMS[assign][c], t)."""
    n = len(donor)
    pos = {(donor[i], codes_of(pheno)[i], tp[i]): i for i in range(n)}
    codes = codes_of(pheno)
    src = np.arange(n)
    for di, d in enumerate(donors_order):
        P = PERMS[assign[di]]
        mask = np.where(donor == d)[0]
        for i in mask:
            src[i] = pos[(d, P[codes[i]], tp[i])]
    return src


def pval(perm_stats, observed):
    b = int(np.sum(np.asarray(perm_stats) >= observed - TOL))
    return (b + 1) / (B + 1)


def bh(pvals):
    p = np.asarray(pvals, dtype=float)
    m = len(p)
    order = np.argsort(p)
    ranked = p[order] * m / np.arange(1, m + 1)
    q = np.minimum.accumulate(ranked[::-1])[::-1]
    out = np.empty(m)
    out[order] = np.clip(q, 0, 1)
    return out


# ---------- core block test on a fixed design family ----------

def _f(ss_block, sse_full, df_block, df_resid):
    return (ss_block / df_block) / (sse_full / df_resid)


def execute(root, write=True):
    tbl = os.path.join(root, "reports", "tables")
    inp = os.path.join(tbl, "PXD016245_time_aware_analysis_input.tsv")
    donor, pheno, tp, Y, rows = load_input(inp)
    codes_obs = codes_of(pheno)
    fixed, time_d = fixed_blocks(donor, tp)

    n = len(donor)
    SST_col = np.sum((Y - Y.mean(axis=0)) ** 2, axis=0)
    SST = float(np.sum(SST_col))

    ZR = fixed
    ZF = build_full(fixed, time_d, codes_obs)
    ZA = build_additive(fixed, codes_obs)
    rank_R = int(np.linalg.matrix_rank(ZR))
    rank_F = int(np.linalg.matrix_rank(ZF))
    rank_A = int(np.linalg.matrix_rank(ZA))
    df_block = rank_F - rank_R
    df_resid = n - rank_F

    design_ok = (rank_R == 8 and rank_F == 23 and rank_A == 11
                 and df_block == 15 and df_resid == 57)

    resid_R, sseR = sse(ZR, Y)
    sseR_col = colsumsq(resid_R)
    resid_F, sseF = sse(ZF, Y)
    sseF_col = colsumsq(resid_F)

    SSb = sseR - sseF
    F_obs = _f(SSb, sseF, 15, 57)
    dR2_obs = SSb / SST

    # per-module and LOMO observed
    F_mod_obs = np.array([_f(sseR_col[m] - sseF_col[m], sseF_col[m], 15, 57) for m in range(8)])
    dR2_mod_obs = np.array([(sseR_col[m] - sseF_col[m]) / SST_col[m] for m in range(8)])
    full7 = sseF - sseF_col
    red7 = sseR - sseR_col
    SST7 = SST - SST_col
    F_lomo_obs = np.array([_f(red7[m] - full7[m], full7[m], 15, 57) for m in range(8)])
    dR2_lomo_obs = np.array([(red7[m] - full7[m]) / SST7[m] for m in range(8)])

    # scaling sensitivity Y
    sd = Y.std(axis=0, ddof=1)
    scaling_blocked = bool(np.any(sd == 0))
    Ys = (Y - Y.mean(axis=0)) / sd if not scaling_blocked else None
    if not scaling_blocked:
        SSTs = float(np.sum((Ys - Ys.mean(axis=0)) ** 2))
        _, sseRs = sse(ZR, Ys)
        _, sseFs = sse(ZF, Ys)
        F_scal_obs = _f(sseRs - sseFs, sseFs, 15, 57)
        dR2_scal_obs = (sseRs - sseFs) / SSTs

    # interaction observed (full vs additive)
    resid_A, sseA = sse(ZA, Y)
    Yhat_A = Y - resid_A
    F_int_obs = _f(sseA - sseF, sseF, 12, 57)
    dR2_int_obs = (sseA - sseF) / SST

    # ---- primary permutation scheme (shared by primary, module, LOMO, scaling) ----
    rng = np.random.Generator(np.random.PCG64(SEED))
    assigns = sample_global(rng, 4, B)
    F_perm = np.empty(B)
    dR2_perm = np.empty(B)
    F_mod_perm = np.empty((8, B))
    F_lomo_perm = np.empty((8, B))
    F_scal_perm = np.empty(B)
    reg_rows = []
    for k, a in enumerate(assigns):
        cperm = permuted_codes(codes_obs, donor, DONORS, a)
        ZFk = build_full(fixed, time_d, cperm)
        rF, ssef = sse(ZFk, Y)
        ssef_col = colsumsq(rF)
        ssb = sseR - ssef
        F_perm[k] = _f(ssb, ssef, 15, 57)
        dR2_perm[k] = ssb / SST
        F_mod_perm[:, k] = [_f(sseR_col[m] - ssef_col[m], ssef_col[m], 15, 57) for m in range(8)]
        f7 = ssef - ssef_col
        F_lomo_perm[:, k] = [_f(red7[m] - f7[m], f7[m], 15, 57) for m in range(8)]
        if not scaling_blocked:
            _, ssefs = sse(ZFk, Ys)
            F_scal_perm[k] = _f(sseRs - ssefs, ssefs, 15, 57)
        reg_rows.append((k + 1, a[0], a[1], a[2], a[3], F_perm[k], dR2_perm[k]))

    p_primary = pval(F_perm, F_obs)
    p_mod = np.array([pval(F_mod_perm[m], F_mod_obs[m]) for m in range(8)])
    q_mod = bh(p_mod)
    p_lomo = np.array([pval(F_lomo_perm[m], F_lomo_obs[m]) for m in range(8)])
    p_scal = pval(F_scal_perm, F_scal_obs) if not scaling_blocked else None

    primary_supported = bool(p_primary <= 0.05 and dR2_obs > 0.02)

    # ---- interaction Freedman-Lane (reuse the same 9999 global assignments) ----
    F_int_perm = np.empty(B)
    for k, a in enumerate(assigns):
        src = residual_src(donor, pheno, tp, DONORS, a)
        Ystar = Yhat_A + resid_A[src]
        _, ssea = sse(ZA, Ystar)
        _, ssef = sse(ZF, Ystar)   # ZF uses observed codes; design fixed, Y* varies
        F_int_perm[k] = _f(ssea - ssef, ssef, 12, 57)
    p_int = pval(F_int_perm, F_int_obs)
    interaction_annot = "INTERACTION_SENSITIVITY_PRESENT" if p_int <= 0.05 else "INTERACTION_SENSITIVITY_ABSENT"

    # ---- leave-one-donor-out ----
    lodo = []
    for d_excl in DONORS:
        keep = donor != d_excl
        dd, pp, tt, YY = donor[keep], pheno[keep], tp[keep], Y[keep]
        cc = codes_of(pp)
        remaining = [d for d in DONORS if d != d_excl]
        fx = np.hstack([np.ones((len(dd), 1)), _dummies(dd, remaining), _dummies(tt, TIMEPOINTS)])
        td = _dummies(tt, TIMEPOINTS)
        ZRk = fx
        ZFk = build_full(fx, td, cc)
        rk_R = int(np.linalg.matrix_rank(ZRk))
        rk_F = int(np.linalg.matrix_rank(ZFk))
        nlo = len(dd)
        ok = (rk_R == 7 and rk_F == 22 and (rk_F - rk_R) == 15 and (nlo - rk_F) == 38)
        if not ok:
            lodo.append(dict(donor_excluded=d_excl, n=nlo, rank_reduced=rk_R, rank_full=rk_F,
                             F="NA", p="NA", delta_R2="NA", joint_criterion="NA",
                             status="SENSITIVITY_BLOCKED"))
            continue
        _, sR = sse(ZRk, YY)
        _, sF = sse(ZFk, YY)
        SSTk = float(np.sum((YY - YY.mean(axis=0)) ** 2))
        Fo = _f(sR - sF, sF, 15, 38)
        dR2o = (sR - sF) / SSTk
        rloc = np.random.Generator(np.random.PCG64(SEED))
        a3 = sample_global(rloc, 3, B)
        Fp = np.empty(B)
        for j, a in enumerate(a3):
            cperm = permuted_codes(cc, dd, remaining, a)
            ZFp = build_full(fx, td, cperm)
            _, sf = sse(ZFp, YY)
            Fp[j] = _f(sR - sf, sf, 15, 38)
        pj = pval(Fp, Fo)
        joint = bool(pj <= 0.05 and dR2o > 0.02)
        lodo.append(dict(donor_excluded=d_excl, n=nlo, rank_reduced=rk_R, rank_full=rk_F,
                         F=Fo, p=pj, delta_R2=dR2o, joint_criterion=joint, status="ok"))

    lodo_all_ok = all(l["status"] == "ok" for l in lodo)
    lodo_all_keep = lodo_all_ok and all(l["joint_criterion"] is True for l in lodo)

    # ---- day-specific (per timepoint), BH across 5 days ----
    day = []
    for t in TIMEPOINTS:
        sel = tp == t
        dd, pp, YY = donor[sel], pheno[sel], Y[sel]
        cc = codes_of(pp)
        fx = np.hstack([np.ones((len(dd), 1)), _dummies(dd, DONORS)])
        ZRk = fx
        ZFk = np.hstack([fx, pheno_onehot(cc)])
        dfb = int(np.linalg.matrix_rank(ZFk)) - int(np.linalg.matrix_rank(ZRk))
        dfr = len(dd) - int(np.linalg.matrix_rank(ZFk))
        _, sR = sse(ZRk, YY)
        _, sF = sse(ZFk, YY)
        SSTk = float(np.sum((YY - YY.mean(axis=0)) ** 2))
        Fo = _f(sR - sF, sF, dfb, dfr)
        dR2o = (sR - sF) / SSTk
        rloc = np.random.Generator(np.random.PCG64(SEED))
        a4 = sample_global(rloc, 4, B)
        Fp = np.empty(B)
        for j, a in enumerate(a4):
            cperm = permuted_codes(cc, dd, DONORS, a)
            ZFp = np.hstack([fx, pheno_onehot(cperm)])
            _, sf = sse(ZFp, YY)
            Fp[j] = _f(sR - sf, sf, dfb, dfr)
        day.append(dict(timepoint=t, F=Fo, p=pval(Fp, Fo), delta_R2=dR2o,
                        df_block=dfb, df_resid=dfr))
    day_q = bh([d["p"] for d in day])
    for d, q in zip(day, day_q):
        d["q_bh"] = float(q)
    d10 = [d for d in day if d["timepoint"] == 10][0]
    day10_annot = ("DAY10_SECONDARY_SIGNAL_PRESENT"
                   if (d10["q_bh"] <= 0.05 and d10["delta_R2"] > 0.02)
                   else "DAY10_SECONDARY_SIGNAL_ABSENT")

    # ---- final decision (four mutually exclusive outcomes) ----
    if not design_ok:
        decision = "EXECUTION_BLOCKED_INVALID_DESIGN"
    elif not primary_supported:
        decision = "ORIGINAL_SEPARATION_NOT_SUPPORTED_AFTER_TIME_ADJUSTMENT"
    elif not lodo_all_keep:
        decision = "AMBIGUOUS_LOW_DONOR_STABILITY_C1"
    else:
        decision = "TIME_ADJUSTED_PHENOTYPE_ARCHITECTURE_SUPPORTED_C1"

    # blocked sensitivities (faithful to prereg: underspecified -> BLOCKED)
    blocked = [
        ("resolution_efferocytosis_LRP1_TGFB1_without_APOE", "SENSITIVITY_BLOCKED",
         "requires recomputing a module score from proteins; the locked input has only "
         "aggregated module scores and rescoring is prohibited"),
        ("restricted_PERMDISP", "SENSITIVITY_BLOCKED",
         "exact restricted-permutation dispersion algorithm is not unambiguously defined "
         "in the sealed prereg; not substituted"),
    ]

    res = dict(
        decision=decision, design_ok=design_ok,
        rank_R=rank_R, rank_F=rank_F, rank_A=rank_A, df_block=df_block, df_resid=df_resid,
        F_obs=F_obs, p_primary=p_primary, dR2_obs=dR2_obs, primary_supported=primary_supported,
        SST=SST, sseR=sseR, sseF=sseF, SSb=SSb,
        F_int_obs=F_int_obs, p_int=p_int, dR2_int_obs=dR2_int_obs, interaction_annot=interaction_annot,
        day10_annot=day10_annot,
        lodo=lodo, lodo_all_keep=lodo_all_keep,
        day=day, blocked=blocked,
        F_mod_obs=F_mod_obs, p_mod=p_mod, q_mod=q_mod, dR2_mod_obs=dR2_mod_obs,
        F_lomo_obs=F_lomo_obs, p_lomo=p_lomo, dR2_lomo_obs=dR2_lomo_obs,
        scaling_blocked=scaling_blocked,
        F_scal_obs=(None if scaling_blocked else F_scal_obs),
        p_scal=p_scal, dR2_scal_obs=(None if scaling_blocked else dR2_scal_obs),
        reg_rows=reg_rows, n_unique_perms=len({tuple(a) for a in assigns}),
        input_sha256=INPUT_SHA256,
    )
    if write:
        _write_tables(tbl, res)
    return res


# ---------- table writers ----------

def _w(path, header, rows):
    with open(path, "w", encoding="utf-8") as f:
        f.write("\t".join(header) + "\n")
        for r in rows:
            f.write("\t".join("" if x is None else str(x) for x in r) + "\n")


def _f6(x):
    return "NA" if x is None or (isinstance(x, float) and (np.isnan(x))) else f"{float(x):.6f}"


def _write_tables(tbl, r):
    _w(os.path.join(tbl, "PXD016245_time_aware_primary_result.tsv"),
       ["metric", "value"],
       [["rank_reduced", r["rank_R"]], ["rank_full", r["rank_F"]], ["rank_additive", r["rank_A"]],
        ["df_block", r["df_block"]], ["df_resid", r["df_resid"]],
        ["SST", _f6(r["SST"])], ["SSE_reduced", _f6(r["sseR"])], ["SSE_full", _f6(r["sseF"])],
        ["SS_block", _f6(r["SSb"])], ["F_observed", _f6(r["F_obs"])],
        ["p_primary", _f6(r["p_primary"])], ["delta_R2", _f6(r["dR2_obs"])],
        ["criterion_p_le_0.05", str(r["p_primary"] <= 0.05)],
        ["criterion_effect_gt_0.02", str(r["dR2_obs"] > 0.02)],
        ["PRIMARY_SUPPORTED", str(r["primary_supported"])],
        ["B", B], ["seed", SEED], ["n_unique_permutations", r["n_unique_perms"]],
        ["input_sha256", r["input_sha256"]]])

    _w(os.path.join(tbl, "PXD016245_time_aware_permutation_registry.tsv"),
       ["permutation_id", "assign_A", "assign_B", "assign_C", "assign_P", "seed",
        "canonical_order", "unique", "F_perm", "delta_R2_perm"],
       [[pid, a, b, c, d, SEED, "M1,M2,dendritic_cell,osteoclast", "true", _f6(F), _f6(dr)]
        for (pid, a, b, c, d, F, dr) in r["reg_rows"]])

    _w(os.path.join(tbl, "PXD016245_time_aware_lodo.tsv"),
       ["donor_excluded", "n", "rank_reduced", "rank_full", "df_block", "df_resid",
        "F", "p", "delta_R2", "joint_criterion", "status"],
       [[l["donor_excluded"], l["n"], l["rank_reduced"], l["rank_full"], 15, 38,
         _f6(l["F"]) if l["F"] != "NA" else "NA", _f6(l["p"]) if l["p"] != "NA" else "NA",
         _f6(l["delta_R2"]) if l["delta_R2"] != "NA" else "NA",
         l["joint_criterion"], l["status"]] for l in r["lodo"]])

    _w(os.path.join(tbl, "PXD016245_time_aware_interaction_sensitivity.tsv"),
       ["test", "df_block", "df_resid", "F_observed", "p", "delta_R2", "annotation", "role"],
       [["interaction_block_full_vs_additive", 12, 57, _f6(r["F_int_obs"]), _f6(r["p_int"]),
         _f6(r["dR2_int_obs"]), r["interaction_annot"],
         "secondary_non_primary_non_decisional (Freedman-Lane residual permutation)"]])

    _w(os.path.join(tbl, "PXD016245_time_aware_day_specific.tsv"),
       ["timepoint", "df_block", "df_resid", "F", "p", "q_bh", "delta_R2", "role"],
       [[d["timepoint"], d["df_block"], d["df_resid"], _f6(d["F"]), _f6(d["p"]),
         _f6(d["q_bh"]), _f6(d["delta_R2"]),
         ("day10_secondary_terminal_time_sensitivity" if d["timepoint"] == 10 else "per_day_BH_family")]
        for d in r["day"]])

    _w(os.path.join(tbl, "PXD016245_time_aware_module_specific.tsv"),
       ["module", "F", "p", "q_bh", "delta_R2", "role"],
       [[MODULES[m], _f6(r["F_mod_obs"][m]), _f6(r["p_mod"][m]), _f6(r["q_mod"][m]),
         _f6(r["dR2_mod_obs"][m]), "descriptive_no_pathway_no_flux_no_mechanism"]
        for m in range(8)])

    _w(os.path.join(tbl, "PXD016245_time_aware_leave_one_module_out.tsv"),
       ["module_removed", "delta_R2_full_space", "delta_R2_without_module",
        "change_in_delta_R2", "F_without_module", "p_without_module", "role"],
       [[MODULES[m], _f6(r["dR2_obs"]), _f6(r["dR2_lomo_obs"][m]),
         _f6(r["dR2_obs"] - r["dR2_lomo_obs"][m]), _f6(r["F_lomo_obs"][m]),
         _f6(r["p_lomo"][m]), "descriptive_only_no_driver_claim"] for m in range(8)])

    _w(os.path.join(tbl, "PXD016245_time_aware_scaling_sensitivity.tsv"),
       ["analysis", "F", "p", "delta_R2", "status", "role"],
       [["unit_variance_module_scaling",
         _f6(r["F_scal_obs"]), _f6(r["p_scal"]), _f6(r["dR2_scal_obs"]),
         ("SENSITIVITY_BLOCKED" if r["scaling_blocked"] else "ok"),
         "supportive_non_decisional"]])

    _w(os.path.join(tbl, "PXD016245_time_aware_resolution_sensitivity.tsv"),
       ["analysis", "delta_R2", "F", "p", "status", "note"],
       [["primary_space_with_resolution_3of6", _f6(r["dR2_obs"]), _f6(r["F_obs"]),
         _f6(r["p_primary"]), "ok", "resolution_efferocytosis covered 3/6 (LRP1,APOE,TGFB1); APOE shared with lipid_foam_cell; not functional efferocytosis"],
        ["exclude_resolution_efferocytosis", _f6(r["dR2_lomo_obs"][5]), _f6(r["F_lomo_obs"][5]),
         _f6(r["p_lomo"][5]), "ok", "7-module space dropping resolution_efferocytosis"],
        ["LRP1_TGFB1_without_APOE", "NA", "NA", "NA", "SENSITIVITY_BLOCKED",
         "requires protein-level rescoring not available from the locked aggregated input"]])

    _w(os.path.join(tbl, "PXD016245_time_aware_blocked_sensitivities.tsv"),
       ["sensitivity", "status", "reason"],
       [[name, status, reason] for (name, status, reason) in r["blocked"]])

    _w(os.path.join(tbl, "PXD016245_time_aware_final_decision.tsv"),
       ["field", "value"],
       [["decision", r["decision"]], ["primary_supported", str(r["primary_supported"])],
        ["p_primary", _f6(r["p_primary"])], ["delta_R2", _f6(r["dR2_obs"])],
        ["leave_one_donor_out_all_keep", str(r["lodo_all_keep"])],
        ["interaction_annotation", r["interaction_annot"]],
        ["day10_annotation", r["day10_annot"]],
        ["claim_ceiling", "C1 descriptive within-dataset time-adjusted"]])

    _w(os.path.join(tbl, "PXD016245_time_aware_claim_trace.tsv"),
       ["claim", "permitted", "basis"],
       [["within-dataset time-adjusted phenotype association (descriptive, C1)", "yes",
         "primary endpoint + prereg claim ceiling"],
        ["mechanism / causality / driver", "no", "blocked by claim ceiling"],
        ["pathway activation / metabolic flux / function", "no", "module scores are abundance summaries"],
        ["validation / integration / convergence / proteome-metabolome coupling", "no", "blocked"],
        ["therapeutic target / clinical utility / population generalization", "no", "blocked"],
        ["C2 / C3", "no", "blocked"]])


def software_environment(sealed_commit, exec_commit):
    import platform
    import sys
    try:
        import pandas as pd
        pdv = pd.__version__
    except Exception:
        pdv = "not_used"
    try:
        import scipy
        spv = scipy.__version__
    except Exception:
        spv = "not_installed"
    return dict(
        python=sys.version.split()[0], numpy=np.__version__, pandas=pdv, scipy=spv,
        os=platform.platform(), sealed_commit=sealed_commit, exec_commit=exec_commit,
        input_sha256=INPUT_SHA256, seed=SEED, B=B,
        utc=datetime.now(timezone.utc).strftime("%Y-%m-%dT%H:%M:%SZ"),
    )
```
