## Supplementary Data 3 - Reproducibility for "Longitudinal proteomic module profiles vary across human monocyte differentiation and polarization conditions": HISTORICAL_SOFTWARE_ENVIRONMENT.md

# PXD016245_TIME_AWARE_SOFTWARE_ENVIRONMENT

- python: 3.10.20
- numpy: 2.2.6
- pandas: 2.3.3
- scipy: 1.15.3
- os: Windows-10-10.0.26200-SP0
- sealed_commit: 3359f629fd3b42dee4719d3a2a74bec151b5a05f
- exec_commit: 3359f629fd3b42dee4719d3a2a74bec151b5a05f
- input_sha256: 1fb757edd9bbb8e2354547ab733fc04a7888cad661b19e488a6e13c4664a8e66
- seed: 20260615
- B: 9999
- utc: 2026-06-19T14:46:25Z
