## Supplementary Data 3 - Reproducibility for "Longitudinal proteomic module profiles vary across human monocyte differentiation and polarization conditions": modules.yml

```
disabled_indices:
  persistence_index:
    status: disabled
    reason: Requires basal, perturbation, recovery/rest, multiple timepoints, same biological system, and sufficient metadata.
  priming_index:
    status: disabled
    reason: Requires basal, perturbation, recovery/rest, multiple timepoints, same biological system, and sufficient metadata.
modules:
  - id: glycolysis_ppp
    description: Glucose uptake, glycolysis, pentose phosphate pathway, NADPH generation.
    proteins:
      - {feature: SLC2A1, direction: positive}
      - {feature: HK2, direction: positive}
      - {feature: PFKP, direction: positive}
      - {feature: ALDOA, direction: positive}
      - {feature: GAPDH, direction: positive}
      - {feature: PKM, direction: positive}
      - {feature: LDHA, direction: positive}
      - {feature: PDK1, direction: positive}
      - {feature: G6PD, direction: positive}
      - {feature: PGD, direction: positive}
    metabolites:
      - {feature: glucose-6-phosphate, direction: positive}
      - {feature: fructose-1,6-bisphosphate, direction: positive}
      - {feature: pyruvate, direction: positive}
      - {feature: lactate, direction: positive}
      - {feature: ribose-5-phosphate, direction: positive}
      - {feature: NADPH, direction: positive}
    functions:
      - {feature: ECAR, direction: positive}
      - {feature: lactate_release, direction: positive}
  - id: mitochondrial_oxphos
    description: Mitochondrial respiration, TCA, OXPHOS, spare respiratory capacity.
    proteins:
      - {feature: PDHA1, direction: positive}
      - {feature: DLAT, direction: positive}
      - {feature: CS, direction: positive}
      - {feature: IDH3A, direction: positive}
      - {feature: OGDH, direction: positive}
      - {feature: SDHA, direction: positive}
      - {feature: MDH2, direction: positive}
      - {feature: NDUFS1, direction: positive}
      - {feature: UQCRC2, direction: positive}
      - {feature: COX4I1, direction: positive}
      - {feature: ATP5F1A, direction: positive}
    metabolites:
      - {feature: citrate, direction: positive}
      - {feature: alpha-ketoglutarate, direction: positive}
      - {feature: succinate, direction: positive}
      - {feature: fumarate, direction: positive}
      - {feature: malate, direction: positive}
    functions:
      - {feature: OCR, direction: positive}
      - {feature: spare_respiratory_capacity, direction: positive}
  - id: lipid_foam_cell
    description: oxLDL uptake, lipid storage, foam-cell-like phenotype.
    proteins:
      - {feature: CD36, direction: positive}
      - {feature: MSR1, direction: positive}
      - {feature: OLR1, direction: positive}
      - {feature: PLIN2, direction: positive}
      - {feature: ACSL1, direction: positive}
      - {feature: SOAT1, direction: positive}
      - {feature: ABCA1, direction: negative}
      - {feature: ABCG1, direction: negative}
      - {feature: APOE, direction: negative}
    metabolites:
      - {feature: cholesteryl_esters, direction: positive}
      - {feature: ceramides, direction: positive}
      - {feature: lysophosphatidylcholines, direction: positive}
      - {feature: acylcarnitines, direction: positive}
    functions:
      - {feature: Dil_oxLDL_uptake, direction: positive}
      - {feature: BODIPY_lipid_droplets, direction: positive}
  - id: redox_ros
    description: NADPH, NOX2, mitochondrial ROS, antioxidant buffering.
    proteins:
      - {feature: CYBB, direction: positive}
      - {feature: NCF1, direction: positive}
      - {feature: NCF2, direction: positive}
      - {feature: NCF4, direction: positive}
      - {feature: SOD2, direction: negative}
      - {feature: GPX1, direction: negative}
      - {feature: PRDX1, direction: negative}
      - {feature: PRDX3, direction: negative}
      - {feature: CAT, direction: negative}
    metabolites:
      - {feature: NADPH, direction: positive}
      - {feature: oxidized_glutathione, direction: positive}
      - {feature: reduced_glutathione, direction: negative}
    functions:
      - {feature: MitoSOX, direction: positive}
      - {feature: DCFDA_ROS, direction: positive}
  - id: inflammasome_cytokine
    description: Inflammasome activation and inflammatory cytokine output.
    proteins:
      - {feature: NLRP3, direction: positive}
      - {feature: PYCARD, direction: positive}
      - {feature: CASP1, direction: positive}
      - {feature: GSDMD, direction: positive}
      - {feature: IL1B, direction: positive}
      - {feature: IL18, direction: positive}
      - {feature: TNF, direction: positive}
      - {feature: IL6, direction: positive}
      - {feature: CXCL8, direction: positive}
    metabolites:
      - {feature: succinate, direction: positive}
      - {feature: lactate, direction: positive}
    functions:
      - {feature: IL1B_release, direction: positive}
      - {feature: IL18_release, direction: positive}
      - {feature: caspase1_activity, direction: positive}
  - id: resolution_efferocytosis
    description: Resolution, efferocytosis, anti-inflammatory and repair state.
    proteins:
      - {feature: MERTK, direction: positive}
      - {feature: AXL, direction: positive}
      - {feature: LRP1, direction: positive}
      - {feature: APOE, direction: positive}
      - {feature: IL10, direction: positive}
      - {feature: TGFB1, direction: positive}
    metabolites:
      - {feature: specialized_proresolving_mediators, direction: positive}
    functions:
      - {feature: efferocytosis_index, direction: positive}
      - {feature: persistent_IL1B, direction: negative}
  - id: monocyte_to_macrophage_differentiation
    description: Normal differentiation background from monocyte to macrophage-like states.
    use: Separate normal differentiation from pathological or stimulus-induced reprogramming.
    proteins:
      - {feature: CSF1R, direction: positive}
      - {feature: CD14, direction: positive}
      - {feature: CD68, direction: positive}
      - {feature: ITGAM, direction: positive}
      - {feature: MRC1, direction: positive}
      - {feature: MARCO, direction: positive}
      - {feature: LYZ, direction: positive}
      - {feature: CTSS, direction: positive}
      - {feature: LST1, direction: positive}
      - {feature: TYROBP, direction: positive}
      - {feature: SPI1, direction: positive}
      - {feature: MAFB, direction: positive}
    metabolites: []
    functions: []
  - id: proteostasis_er_stress
    description: Protein folding, ER stress, unfolded protein response, lipotoxic stress adaptation.
    use: Connect lipotoxicity, ROS, inflammation, and cellular stress.
    proteins:
      - {feature: HSPA5, direction: positive}
      - {feature: HSP90B1, direction: positive}
      - {feature: CALR, direction: positive}
      - {feature: CANX, direction: positive}
      - {feature: PDIA3, direction: positive}
      - {feature: PDIA4, direction: positive}
      - {feature: XBP1, direction: positive}
      - {feature: ATF4, direction: positive}
      - {feature: DDIT3, direction: positive}
      - {feature: SQSTM1, direction: positive}
    metabolites: []
    functions: []
```
