## Supplementary Data 3 - Reproducibility for "Longitudinal proteomic module profiles vary across human monocyte differentiation and polarization conditions": PXD016245_STATE_SEPARATION_TIME_AWARE_PREREG.md

# PXD016245 time-aware state-separation pre-registration (amendment A001, revision R5)

> STATUS: **SEALED (A001-R5) — 2026-06-19.** Sealed by explicit human action; the
> SHA-256 of this sealed document is recorded in `registry/PREREG_HASHES.md`. This file
> is now frozen and read-only; any further change requires a dated amendment in
> `00_prereg/AMENDMENTS.md`. Confirmatory execution is authorized **only** on a dedicated
> analysis branch created from the sealing commit. No analysis was run under this plan
> before sealing.
>
> Revision history (this document supersedes the prior revisions; it is **R5**):
> - **R1** (2026-06-19): time-aware model and reduced-vs-full joint-block endpoint.
> - **R2** (2026-06-19): trajectory-preserving primary permutation.
> - **R3** (2026-06-19): lock the longitudinal analytical input and its hashes, lock
>   the scoring provenance, replace the per-module mixed model with a fixed-effect
>   block test, fix the permutation p-value algorithm, and make the decision
>   categories a mutually exclusive tree.
> - **R4** (2026-06-19): final statistical closure before sealing — unify the design
>   variable name to `phenotype` (the real input column), close the exact pseudo-F /
>   rank / R2 definitions, fix the permutation generator, demote day-10 to a secondary
>   terminal-time sensitivity inside the five-day BH family, reduce the decision tree to
>   five mutually exclusive outcomes, and add a non-decisional unit-variance module
>   scaling sensitivity.
> - **R5** (2026-06-19): remove the interaction test from the decision tree. Direct
>   permutation of `phenotype` labels destroys both the phenotype main effect and the
>   interaction simultaneously, so a "full vs additive" test under that null tests
>   "no phenotype effect at all", not "a constant phenotype effect with no interaction";
>   it therefore cannot be a decisional split. The decision tree is reduced to four
>   outcomes (dropping `TIME_DEPENDENT_TRAJECTORY_DIFFERENCE_C1`); the interaction is kept
>   only as a non-decisional secondary sensitivity with an exact Freedman-Lane residual
>   permutation under the additive model. The solver is fixed to
>   `numpy.linalg.lstsq(Z, Y, rcond=None)` and leave-one-donor-out expected ranks are
>   stated. No analysis executed; no result observed; changes made before sealing.

## 1. Motivation
The original analysis pooled five differentiation time points (days 2,4,6,8,10) per
donor x phenotype as replicates and blocked permutations on donor only
(see `PXD016245_LONGITUDINAL_DESIGN_AUDIT.md`). This plan tests whether differentiation
phenotype — including phenotype-specific temporal trajectories — explains variation
beyond donor and time, on an explicitly materialized longitudinal input.

## 2. Hypothesis (to test, not to confirm)
After adjusting for donor and differentiation time, the differentiation phenotype
(including phenotype-specific temporal trajectories) explains additional variation in
module-score space within PXD016245. A negative result is acceptable and publishable.

## 3. Design and units
- Independent biological units: **n=4 donors** (A, B, C, P), repeated measures.
- **80 repeated longitudinal observations** (not 80 independent samples).
- Conditions (the `phenotype` column; see §3a): dendritic_cell, M1, M2, osteoclast.
- Time: differentiation days 2, 4, 6, 8, 10 (categorical primary; see §4).
- Baseline monocyte (one per donor, day 0) excluded from the primary input.
- Power and generalizability are limited by n=4 donors; any result remains a
  descriptive within-dataset inference (claim ceiling: C1 descriptive).

## 3a. Design variable name (phenotype)
The design variable is named **`phenotype`** in every model, endpoint, table and test of
this revision, because `phenotype` is the actual column name of the locked input
`reports/tables/PXD016245_time_aware_analysis_input.tsv`. It corresponds exactly to the
historical label `cell_identity`; its values are copied without transformation
(`phenotype := cell_identity`, an identity copy). `phenotype` is an **operational label**
of the four monocyte-derived differentiation/polarization conditions and is **not** an
ontological assertion that M1, M2, dendritic_cell and osteoclast are equivalent lineages.
`cell_identity` is retained only as the historical name.

## 4. Time coding
Primary analysis uses time as a categorical factor `factor(timepoint)` with five
levels (2,4,6,8,10). Numeric-linear time is a sensitivity only.

## 5. Locked analytical input (materialized, hash-locked)
- **Input file:** `reports/tables/PXD016245_time_aware_analysis_input.tsv`
- **SHA-256:** `1fb757edd9bbb8e2354547ab733fc04a7888cad661b19e488a6e13c4664a8e66`
- Shape: **80 rows** x (5 metadata columns + 8 module-score columns).
- Columns: sample_id, sample_column, donor_id, phenotype, timepoint,
  glycolysis_ppp, mitochondrial_oxphos, lipid_foam_cell, redox_ros,
  inflammasome_cytokine, resolution_efferocytosis,
  monocyte_to_macrophage_differentiation, proteostasis_er_stress.
- **Time materialization rule:** suffixes `_2/_4/_6/_8/_10` -> timepoint 2/4/6/8/10;
  `Mono` samples -> day 0, excluded from the primary input; the historical
  `timepoint=unknown` column is not used; the historical `replicate` column is
  recognized as a mislabeled time label, not a replicate.
- **Validation (recorded in `PXD016245_time_aware_input_qc.tsv`):** 4 donors x 4
  phenotypes x 5 timepoints = 80 rows; no `unknown`; no duplicate or missing
  donor x phenotype x timepoint combination; eight numeric scores per row; no
  monocytes in the primary input.
- Companion manifest with hashes of the input, the frozen score table, module
  definitions, the source quantitative matrix and the scoring script:
  `reports/tables/PXD016245_time_aware_input_manifest.tsv`.
- The locked input must not be modified by this revision; its SHA-256 above must remain
  unchanged.

## 5b. Scoring provenance lock (no rescoring during A001)
The module scores in the input were computed previously and are **frozen**. They were
derived by: (1) using positive LFQ intensities (non-numeric, zero and negative values
treated as missing); (2) log2 transformation; (3) standardizing each protein across
**all 84 samples** (including the 4 day-0 monocytes) using observed values only, with the
sample standard deviation (ddof=1) and no imputation; (4) orienting negative-direction
proteins; (5) averaging the available oriented per-protein z-scores per module (arithmetic
mean), requiring at least two usable proteins per module per sample and a per-protein
missingness <= 0.5. **Rescoring, re-standardization, or recomputation of module scores
during A001 is prohibited.** In particular, proteins were standardized over the 84-sample
scale even though the primary test uses 80 observations; this scale is fixed and must not
be recomputed over the 80-observation subset.

## 6. Models (donor explicit, fixed effects only)
Let `Y` be the locked 80x8 numeric matrix of the eight module scores.
```
Reduced:  Y ~ donor + factor(timepoint)
Full:     Y ~ donor + factor(timepoint) + phenotype + factor(timepoint):phenotype
Additive: Y ~ donor + factor(timepoint) + phenotype
```
Donor is a **fixed effect** in the model (no random intercept; with only four donors a
random effect is unstable) and is separately preserved by the permutation restriction.

## 7. Primary endpoint (single, block comparison)
Joint phenotype block `phenotype + factor(timepoint):phenotype`, evaluated as a
reduced-vs-full comparison (marginal/incremental, not order-dependent sequential):
- permutation p-value for the joint block;
- incremental R2 = R2(full) - R2(reduced);
- pseudo-F from the reduced-versus-full sums of squares (§7a). This multivariate linear
  model on the eight-module matrix is equivalent to the Euclidean distance-based PERMANOVA
  partition, and is defined so the algorithm is unambiguous.

## 7a. Exact multivariate statistic (locked)
Definitions:
- `Y`: the numeric **80x8** matrix of the eight module scores from the locked input.
- `Z_R`: design matrix of the reduced model; `Z_F`: design matrix of the full model.
- All models **include an intercept**. Factors use a **full-rank** representation; the
  inference depends on the column space of the design, not on the choice of reference
  category.
- Fit by least squares using **only** `numpy.linalg.lstsq(Z, Y, rcond=None)` (no `pinv`
  alternative) for coefficients and fitted values; validate ranks with
  `numpy.linalg.matrix_rank`. For each model M:
```
beta_M  = numpy.linalg.lstsq(Z_M, Y, rcond=None)[0]
Y_hat_M = Z_M @ beta_M
SSE_M   = || Y - Y_hat_M ||_F^2      (squared Frobenius norm)
```
- Total sum of squares about the column means:
```
SST       = || Y - column_mean(Y) ||_F^2
SS_block  = SSE_reduced - SSE_full
df_block  = rank(Z_full) - rank(Z_reduced)
df_resid  = n - rank(Z_full)              (n = 80)
```
- Incremental pseudo-F and R2:
```
F_block   = (SS_block / df_block) / (SSE_full / df_resid)
R2_M      = 1 - SSE_M / SST
delta_R2  = R2_full - R2_reduced = SS_block / SST
```
- **Expected design ranks** (validated with `numpy.linalg.matrix_rank` before any
  permutation):
  rank(Z_reduced) = **8**, rank(Z_additive) = **11**, rank(Z_full) = **23**, df_block = **15**, df_resid = **57**.
  Any deviation from rank(Z_reduced)=8 or rank(Z_full)=23 triggers
  `EXECUTION_BLOCKED_INVALID_DESIGN`.
- **Leave-one-donor-out expected ranks** (3 donors, 60 observations):
  rank(Z_reduced) = **7**, rank(Z_full) = **22**, df_block = **15**, df_resid = **38**.
  A deviation triggers the stop rule for that sensitivity and, if the primary endpoint had
  passed, yields `AMBIGUOUS_LOW_DONOR_STABILITY_C1`.

## 8. Primary permutation scheme (trajectory-preserving)
The repeated unit is the `donor x phenotype` trajectory observed at days 2,4,6,8,10.
Within each donor, randomly permute the four phenotype trajectory labels once and then
apply the same permutation consistently across all five timepoints for that donor.
Do not permute timepoint; do not swap observations between donors; do not permute
timepoints independently. Permutation space: 4! = 24 per donor, 24^4 = 331,776 global.

## 8a. Permutation generator (locked)
- **Canonical phenotype order:** `M1, M2, dendritic_cell, osteoclast`. Enumerate the 24
  permutations of these four labels.
- For the primary analysis, the per-donor permutations form the Cartesian product over
  donors A, B, C, P (24^4 = 331,776). **Exclude the identity assignment**, then select
  **9999** distinct combinations **without replacement**.
- RNG: `numpy.random.Generator(numpy.random.PCG64(20260615))`.
- Each selected per-donor permutation is applied **once per donor, identically across the
  five timepoints**, so each trajectory keeps its five-point shape.
- The exact list of permutations actually used is saved as an execution artifact **after
  sealing** (not created by this revision).

## 8b. Permutation algorithm (locked)
- B = **9999** permutations; **seed = 20260615**.
- p-value: **p = (b + 1) / (B + 1)**, where `b` is the number of permuted statistics
  greater than or equal to the observed statistic, using the tie rule
  `F_perm >= F_observed - 1e-9`.
- The observed (identity) assignment is **excluded** from the B permutations and added
  via the `+1` terms.
- Sampling is **without replacement** over distinct permutations; duplicate permutations
  are rejected and redrawn; if the finite space is smaller than B (e.g.
  leave-one-donor-out, 24^3 = 13,824 > 9999, still ample), use 9999 distinct random
  draws and document.
- In each permutation, `phenotype` is reassigned and `Z_F` is rebuilt; `Z_R` is reused
  (it does not depend on phenotype); SSE_reduced, SSE_full, SS_block, incremental R2 and
  pseudo-F are recomputed.
- Implementation: a documented in-repo function (numpy/scipy); library and versions
  recorded in `99_report/env.lock` at execution. Numerical equality tolerance = 1e-9.

## 9. Three nulls (with explicit decision weight)
1. **Primary (decisional):** trajectory-preserving permutation (§8, §8a, §8b).
2. **Known-signal sensitivity (supportive, not decisional):** for each module, fit
   `score ~ donor + factor(timepoint)`, take residuals, and test phenotype on the
   residuals using the **same trajectory-preserving permutation**. `factor(timepoint)`
   is fully flexible across the five days, so no nonlinear smoother is fitted and no
   smoothing parameters or degrees of freedom are chosen after seeing the data.
3. **Free label shuffle (diagnostic only, NON-decisional):** unrestricted permutation
   ignoring strata and trajectories; the result is NOT required to "survive" this null.

### 9b. Timepoint-stratified label permutation sensitivity (supportive, non-primary)
The R1 independent `donor x timepoint` permutation is retained only as a supportive,
non-primary, non-decisional sensitivity; it does not preserve complete phenotype
trajectories.

## 10. Per-module analysis (fixed effects; no mixed model)
Per module, reduced-vs-full block test (NOT a random-effects mixed model):
```
Reduced:  score ~ donor + factor(timepoint)
Full:     score ~ donor + factor(timepoint) + phenotype + factor(timepoint):phenotype
```
- donor as a fixed effect; test the joint `phenotype + interaction` block;
- same trajectory-preserving permutation (9999, seed 20260615, p=(b+1)/(B+1));
- Benjamini-Hochberg correction across the **8 modules**.
No random-intercept term for donor is used or implied anywhere; donor enters only as a
fixed effect (with four donors a random intercept is unstable).

## 11. Pre-specified sensitivities
1. **Day-10 test (pre-specified SECONDARY terminal-time sensitivity):**
   `Y ~ donor + phenotype` at day 10 only; permute phenotype within donor; reported with
   the four other per-day tests under one Benjamini-Hochberg family (§11.2). It is **not**
   a confirmatory test (not independent of the global analysis) and **cannot rescue a
   negative primary endpoint**. Its
   only role is a descriptive annotation `DAY10_SECONDARY_SIGNAL_PRESENT` /
   `DAY10_SECONDARY_SIGNAL_ABSENT`, defined by BH-adjusted p <= 0.05 AND incremental
   R2 (vs `Y ~ donor` at day 10) > 0.02; this annotation does not change the final
   decision (§13).
2. **Per-day tests at days 2,4,6,8,10:** phenotype within donor; the five day-specific
   p-values form **one Benjamini-Hochberg family** (descriptive, non-decisional).
3. Additive model without interaction; trajectory permutation.
4. **Interaction-block secondary sensitivity (supportive, non-primary, non-decisional).**
   The interaction `factor(timepoint):phenotype` is tested **only** as a secondary
   sensitivity that can neither rescue nor invalidate the primary result, because direct
   permutation of `phenotype` labels destroys the phenotype main effect as well as the
   interaction. To isolate the interaction under a valid null we use an exact
   Freedman-Lane residual permutation under the additive model:
   (a) fit the observed additive model `Y ~ donor + factor(timepoint) + phenotype` to get
   `Yhat_additive` and residuals `E_additive = Y - Yhat_additive`;
   (b) permute the residuals as complete trajectories within each donor (one permutation of
   the four trajectory labels per donor, applied identically to the five timepoints; no
   mixing across donors; timepoints not permuted);
   (c) form `Y_star = Yhat_additive + permuted(E_additive)`;
   (d) refit the additive and full models to each `Y_star` and compute the pseudo-F of the
   interaction block `factor(timepoint):phenotype` (df_block = rank(Z_full)-rank(Z_additive)
   = 12);
   (e) B = 9999, seed = 20260615, p = (b + 1) / (B + 1).
   Output is **only** an annotation: `INTERACTION_SENSITIVITY_PRESENT` (permutation
   p <= 0.05; incremental R2 is not a requirement for this annotation),
   `INTERACTION_SENSITIVITY_ABSENT` (p > 0.05), or `INTERACTION_SENSITIVITY_BLOCKED` if the
   exact residual scheme cannot be implemented. This annotation never changes the §13
   decision and never blocks the primary endpoint.
5. Numeric-linear time coding (sensitivity only).
6. Restricted PERMDISP under the trajectory-preserving scheme where applicable; if not
   unambiguously applicable, record as **BLOCKED** (do not substitute another scheme).
7. Module-wise tests (§10) with BH across 8 modules.
8. Time-conditioned leave-one-module-out influence.
9. **Leave-one-donor-out stability (principal stability; replaces bootstrap):** rerun
   the primary omitting A, then B, then C, then P; trajectory permutation within the
   three remaining donors (space 24^3 = 13,824; 9999 distinct random draws; seed
   20260615; expected ranks rank(Z_reduced)=7, rank(Z_full)=22, df_block=15,
   df_resid=38 per §7a); report incremental R2, p, and decision stability for each.
   Bootstrap is not used as the principal stability analysis: with only four donors,
   bootstrap resampling of donors is unstable and no bootstrap confidence intervals are
   constructed.
10. **Unit-variance module scaling (supportive, non-decisional):** the frozen score
    matrix without rescaling is the primary analysis (continuity with the prior plan);
    as a non-decisional sensitivity, center each of the eight modules to its mean over the
    80 locked observations and divide by its sample standard deviation (ddof=1), then rerun
    the same models and permutations and compare direction, p and delta_R2. If any module
    has zero standard deviation over the 80 rows, record this sensitivity as **BLOCKED**.
    This sensitivity cannot by itself rescue or invalidate the primary decision. Rationale:
    the modules have unequal dispersion, so the unscaled Euclidean statistic weights
    higher-dispersion modules (e.g. mitochondrial_oxphos, glycolysis_ppp) more.
11. resolution/efferocytosis sensitivities (module 3/6 covered: LRP1, APOE, TGFB1
    detected; MERTK, AXL, IL10 not detected; APOE shared with lipid_foam_cell):
    - original partial module (3/6);
    - exclusion of the whole module;
    - score excluding APOE, using only LRP1 and TGFB1;
    - none measures functional efferocytosis.

## 12. Primary support threshold
Phenotype architecture is "supported" only if, under the primary trajectory-preserving
permutation: **p <= 0.05 AND incremental R2 > 0.02** for the joint phenotype block. The
0.02 incremental-R2 threshold is a preregistered practical-relevance filter, not a
universal biological cutoff.

## 13. Decision tree (four mutually exclusive outcomes, evaluated in order)
Exactly one code is assigned by walking this tree top to bottom:
1. If a stop rule (§15) fires (including a design-rank mismatch) ->
   `EXECUTION_BLOCKED_INVALID_DESIGN`.
2. Else if the primary does **not** meet the criterion (§12: p <= 0.05 AND
   incremental R2 > 0.02) -> `ORIGINAL_SEPARATION_NOT_SUPPORTED_AFTER_TIME_ADJUSTMENT`.
3. Else if the primary meets the criterion **but** at least one leave-one-donor-out
   re-analysis loses the joint criterion (p>0.05 or incremental R2<=0.02) ->
   `AMBIGUOUS_LOW_DONOR_STABILITY_C1`.
4. Else (primary meets the criterion and all four leave-one-donor-out runs keep it) ->
   `TIME_ADJUSTED_PHENOTYPE_ARCHITECTURE_SUPPORTED_C1`.

"Loses the joint criterion" in step 3 means an objective failure of the §12 threshold in
at least one of the four leave-one-donor-out runs (not a subjective judgement). The
interaction test is **not** a decisional split (the direct-label-permutation null is
invalid for it; see §11.4): the interaction is reported only as the non-decisional
secondary annotation `INTERACTION_SENSITIVITY_PRESENT/ABSENT/BLOCKED`. The day-specific,
module-specific, unit-variance-scaling and interaction results are annotations or
sensitivities; they never substitute for this primary decision. There is no separate
terminal-only or time-dependent decision code: a day-10 signal is reported only as the
secondary annotation in §11.1.

## 14. Claim ceiling
Descriptive within-dataset inference only. A supported result licenses, at most, a
descriptive, time-adjusted within-dataset separation claim. No mechanism, causality,
convergence, integration, validation, flux, trajectory-as-mechanism, therapeutic or
clinical-utility claim.

## 15. Stop rules
- Locked input hash mismatch, missing input, or any `unknown`/duplicate/missing
  combination at execution: `EXECUTION_BLOCKED_INVALID_DESIGN`.
- Design-rank mismatch (rank(Z_reduced) != 8 or rank(Z_full) != 23):
  `EXECUTION_BLOCKED_INVALID_DESIGN`.
- Trajectory-preserving permutation not implementable as specified:
  `EXECUTION_BLOCKED_INVALID_DESIGN`.
- Any rescoring/re-standardization, full-proteome substitution, cross-dataset input,
  multiomic integration, or PCA-as-inference: stop.

## 16. Companion specification tables
- `reports/tables/PXD016245_time_aware_analysis_input.tsv` (locked input)
- `reports/tables/PXD016245_time_aware_input_manifest.tsv` (hashes)
- `reports/tables/PXD016245_time_aware_input_qc.tsv` (validation)
- `reports/tables/PXD016245_time_aware_prereg_models.tsv`
- `reports/tables/PXD016245_time_aware_prereg_endpoints.tsv`
- `reports/tables/PXD016245_time_aware_prereg_permutations.tsv`
- `reports/tables/PXD016245_time_aware_prereg_sensitivities.tsv`
- `reports/tables/PXD016245_time_aware_prereg_decision_rules.tsv`
- `reports/tables/PXD016245_time_aware_prereg_claim_boundaries.tsv`

## 17. Outputs (created only at execution, AFTER sealing)
Reduced/full partition, joint-block p and incremental R2, known-signal sensitivity,
leave-one-donor-out stability, module-wise BH results, time-conditioned module
influence, restricted PERMDISP, unit-variance scaling sensitivity, day-10 secondary
annotation, the interaction-block secondary annotation (Freedman-Lane residual scheme,
§11.4), the saved permutation list, and a single decision code from §13. No result file
is created by this revision.

## 18. Sealing
Sealed on 2026-06-19 by explicit human action. The SHA-256 of this sealed document is
recorded in `registry/PREREG_HASHES.md`. The plan is frozen; any change now requires a
dated amendment in `00_prereg/AMENDMENTS.md`, never an edit here.
**SEALED — confirmatory execution authorized on an analysis branch created from the
sealing commit.**
