## Supplementary Data 3 - Reproducibility for "Longitudinal proteomic module profiles vary across human monocyte differentiation and polarization conditions": README.md

# Supplementary Data 3: primary-result verification from frozen module scores

This compact package independently verifies the primary PXD016245 result reported in the revised manuscript. It starts with the frozen 80-observation by eight-module score matrix and the exact 9,999 archived trajectory-preserving permutation assignments. The new verifier does not import the historical analysis implementation.

Read `DATA_DESIGN_CORRECTION.md` before interpreting biological replication. The archived `donor_id` column contains four source-block labels parsed from filenames. The source study describes three individual-donor preparations and a preparation pooled from 40 donors; these are not four individual donors. The correspondence of the label P to the pooled preparation is inferred, not explicitly verified.

## Run

Use Python 3.10 or later with NumPy and pandas installed. The exact dependency versions used to check this package are in `requirements.txt`; installation, if needed, is `python -m pip install -r requirements.txt`.

From the extracted package directory, run:

```console
python verify_frozen_results.py
```

Expected output begins:

```text
PASS: F=4.257567202; delta_R2=0.313662853; p=0.0002; b=1; B=9999
```

The command exits with code 0 on success or a nonzero code on a failed check. It writes `verification/primary_verification.json`. Its paths are relative to the script, so it can also be launched using an absolute script path. No source or frozen data file is rewritten.

## What is verified

- SHA-256 and byte size of every distributed input, provenance file, source file, environment specification, and verifier listed in `SHA256SUMS.tsv`.
- Historical LF-normalized hashes for the input, preregistration, permutation registry, primary-result table and historical execution module, using selected rows from the original result manifest.
- The preregistration hash against the original project-level registry entry. The original registry records 2026-06-19 and the historical software environment records the sealing/execution commit. These records are preserved as documentary provenance, not independently certified timestamps.
- The complete 4-source-block by 4-condition by 5-day design, 80 unique observations and finite values in all eight modules.
- The reduced, additive and full model ranks (8, 11 and 23) and observed sums of squares, incremental R2 and pseudo-F.
- Every archived assignment, preserving each source-block/condition trajectory across days, with a different reference coding from the original execution. All 9,999 recomputed pseudo-F and incremental-R2 values must agree with the archived six-decimal values to their rounding precision.
- The observed extreme count (1) and permutation p value, `(1+1)/(9999+1)=0.0002`.

The primary endpoint is the joint condition block, including condition and condition-by-time terms, beyond source block and categorical time. It is not a condition main-effect-only test. The four source blocks contribute 80 serial culture observations; neither 80 observations nor the 40 members of a pooled preparation provide that many independent analytical replicates. Numerical verification does not validate the biological independence or population representativeness of these preparations.

## Scope and provenance

This package verifies only the primary endpoint from previously frozen scores. It does not process mass-spectrometry raw files, reconstruct the protein-module scores, rerun leave-one-source-block-out or other secondary analyses, or provide independent biological replication. The historical name of the block-exclusion analysis was leave-one-donor-out. The module dictionary is included to document the original definitions and signs, not as an executable recipe for rescoring in this package. The scores were standardized on the original 84 samples before exclusion of four day-0 monocyte observations.

`frozen/` contains unmodified source input, permutation registry and primary result. `provenance/` contains the original sealed PXD016245 plan, dictionary, selected original registry/manifest entries and software metadata. `historical_source/` contains the original execution and audit modules for inspection only; their broader project dependencies and result inputs are not all distributed here. The documented entrypoint is the new `verify_frozen_results.py`.

Original source: historical COSMOS project, `discovery/cosmos-myeloid-public`. The preregistration entry was extracted unchanged from the registry in the original submission archive. Its plan describes a new time-adjusted analysis after earlier exploration of the same public dataset. It should not be described as analysis of previously unseen data. Original terminology and previous claims in the archived plan/source files are retained for traceability; the revised manuscript governs current interpretation.

The historical Metabolomics Workbench analysis and old manuscripts are excluded. The package manifest excludes itself and the generated verification report to avoid circular hashes and to allow verification under another compatible runtime. Archive-level integrity is separately recorded alongside the ZIP.
