## Supplementary Data 3 - Reproducibility for "Longitudinal proteomic module profiles vary across human monocyte differentiation and polarization conditions": verify_frozen_results.py

```
#!/usr/bin/env python3
"""Independently verify the PXD016245 primary endpoint from frozen scores.

Run: python verify_frozen_results.py
The script reads files relative to itself and writes only verification/primary_verification.json.
It does not import the historical implementation, regenerate module scores, process
raw spectra, regenerate permutations, or verify secondary analyses.
"""
from __future__ import annotations

import csv
import hashlib
import itertools
import json
from pathlib import Path
import platform
import sys

import numpy as np
import pandas as pd

BASE = Path(__file__).resolve().parent
MODULES = [
    "glycolysis_ppp", "mitochondrial_oxphos", "lipid_foam_cell", "redox_ros",
    "inflammasome_cytokine", "resolution_efferocytosis",
    "monocyte_to_macrophage_differentiation", "proteostasis_er_stress",
]
# Legacy schema: these labels identify source blocks, not four individual donors.
# See DATA_DESIGN_CORRECTION.md; retain the numerical coding and frozen column name.
DONORS = ["A", "B", "C", "P"]
CONDITIONS = ["M1", "M2", "dendritic_cell", "osteoclast"]
DAYS = [2, 4, 6, 8, 10]
INPUT_RAW_SHA256 = "1fb757edd9bbb8e2354547ab733fc04a7888cad661b19e488a6e13c4664a8e66"
PREREG_LF_SHA256 = "ce2d30280a7619b1d7c7f74913d6d616e90819682f7025cfe3949038a0d15ef6"
ROUNDING_TOLERANCE = 5.001e-7
PERMUTATION_TIE_TOLERANCE = 1e-9


class VerificationError(RuntimeError):
    pass


def require(condition, message):
    if not bool(condition):
        raise VerificationError(message)


def digest(path, lf_normalized=False):
    data = path.read_bytes()
    if lf_normalized:
        data = data.replace(b"\r\n", b"\n")
    return hashlib.sha256(data).hexdigest()


def verify_package_hashes():
    manifest = BASE / "SHA256SUMS.tsv"
    with manifest.open(encoding="utf-8", newline="") as handle:
        records = list(csv.DictReader(handle, delimiter="\t"))
    seen = set()
    for row in records:
        relative = row["path"]
        path = (BASE / relative).resolve()
        require(path.is_relative_to(BASE), f"Manifest path escapes package: {relative}")
        require(relative not in seen, f"Duplicate manifest entry: {relative}")
        seen.add(relative)
        require(path.is_file(), f"Missing packaged file: {relative}")
        require(path.stat().st_size == int(row["size_bytes"]), f"Size mismatch: {relative}")
        require(digest(path) == row["sha256"], f"SHA-256 mismatch: {relative}")
    required = {
        "verify_frozen_results.py",
        "frozen/PXD016245_time_aware_analysis_input.tsv",
        "frozen/PXD016245_time_aware_permutation_registry.tsv",
        "frozen/PXD016245_time_aware_primary_result.tsv",
        "provenance/PXD016245_STATE_SEPARATION_TIME_AWARE_PREREG.md",
        "provenance/modules.yml",
        "provenance/HISTORICAL_RESULT_MANIFEST_SELECTED_ROWS.tsv",
        "provenance/HISTORICAL_PREREGISTRY_ENTRY.txt",
    }
    require(required <= seen, "Package manifest omits a required verification input")
    return len(records)


def verify_historical_hashes():
    path = BASE / "provenance/HISTORICAL_RESULT_MANIFEST_SELECTED_ROWS.tsv"
    with path.open(encoding="utf-8", newline="") as handle:
        rows = list(csv.DictReader(handle, delimiter="\t"))
    for row in rows:
        target = BASE / row["package_path"]
        require(digest(target, True) == row["sha256_lf_normalized"],
                f"Historical LF-normalized SHA-256 mismatch: {row['package_path']}")
    prereg = BASE / "provenance/PXD016245_STATE_SEPARATION_TIME_AWARE_PREREG.md"
    require(digest(prereg, True) == PREREG_LF_SHA256, "Preregistration differs from recorded seal")
    require(PREREG_LF_SHA256 in (BASE / "provenance/HISTORICAL_PREREGISTRY_ENTRY.txt").read_text(encoding="utf-8"),
            "Historical preregistration entry does not contain the expected seal")
    return len(rows)


def dummies_drop_last(values, categories):
    return np.column_stack([values == category for category in categories[:-1]]).astype(float)


def residual_ss(design, response):
    fitted = design @ np.linalg.lstsq(design, response, rcond=None)[0]
    return float(np.sum((response - fitted) ** 2))


def verify():
    n_hashes = verify_package_hashes()
    n_historical_hashes = verify_historical_hashes()
    input_path = BASE / "frozen/PXD016245_time_aware_analysis_input.tsv"
    require(digest(input_path) == INPUT_RAW_SHA256, "Locked input SHA-256 differs from original")
    frame = pd.read_csv(input_path, sep="\t", dtype={"donor_id": str})
    expected_columns = ["sample_id", "sample_column", "donor_id", "phenotype", "timepoint", *MODULES]
    require(list(frame.columns) == expected_columns, "Unexpected input columns or column order")
    require(len(frame) == 80, "Expected 80 repeated observations")
    require(frame.sample_id.is_unique, "Duplicate sample IDs")
    combinations = list(zip(frame.donor_id, frame.phenotype, frame.timepoint))
    expected_combinations = set(itertools.product(DONORS, CONDITIONS, DAYS))
    require(len(set(combinations)) == 80 and set(combinations) == expected_combinations,
            "Input is not the complete 4-source-block by 4-condition by 5-day design")

    y = frame[MODULES].to_numpy(dtype=float)
    require(np.isfinite(y).all(), "Nonfinite module score")
    donor = frame.donor_id.to_numpy()
    day = frame.timepoint.to_numpy()
    codes = np.array([CONDITIONS.index(label) for label in frame.phenotype], dtype=int)
    donor_codes = np.array([DONORS.index(label) for label in donor], dtype=int)
    time_columns = dummies_drop_last(day, DAYS)
    reduced = np.column_stack([np.ones(80), dummies_drop_last(donor, DONORS), time_columns])

    def full_design(condition_codes):
        phenotype_columns = dummies_drop_last(condition_codes, list(range(4)))
        interactions = (time_columns[:, :, None] * phenotype_columns[:, None, :]).reshape(80, -1)
        return np.column_stack([reduced, phenotype_columns, interactions])

    full = full_design(codes)
    additive = np.column_stack([reduced, dummies_drop_last(codes, list(range(4)))])
    ranks = [int(np.linalg.matrix_rank(design)) for design in (reduced, additive, full)]
    require(ranks == [8, 11, 23], f"Unexpected model ranks: {ranks}")
    df_block, df_residual = ranks[2] - ranks[0], len(y) - ranks[2]
    total_ss = float(np.sum((y - y.mean(axis=0)) ** 2))
    reduced_ss, full_ss = residual_ss(reduced, y), residual_ss(full, y)
    block_ss = reduced_ss - full_ss
    observed_f = (block_ss / df_block) / (full_ss / df_residual)
    delta_r2 = block_ss / total_ss

    archived = pd.read_csv(BASE / "frozen/PXD016245_time_aware_primary_result.tsv", sep="\t")
    require(archived.metric.is_unique, "Duplicated archived primary metric")
    expected = dict(zip(archived.metric, archived.value))
    values = {
        "rank_reduced": ranks[0], "rank_additive": ranks[1], "rank_full": ranks[2],
        "df_block": df_block, "df_resid": df_residual,
        "SST": total_ss, "SSE_reduced": reduced_ss, "SSE_full": full_ss,
        "SS_block": block_ss, "F_observed": observed_f, "delta_R2": delta_r2,
    }
    for metric, value in values.items():
        require(abs(value - float(expected[metric])) <= ROUNDING_TOLERANCE,
                f"Observed statistic disagrees with archived result: {metric}")
    require(expected["input_sha256"] == INPUT_RAW_SHA256, "Archived input hash differs")
    require(int(expected["B"]) == 9999 and int(expected["seed"]) == 20260615,
            "Archived permutation count/seed differs")

    registry = pd.read_csv(BASE / "frozen/PXD016245_time_aware_permutation_registry.tsv", sep="\t")
    require(len(registry) == 9999, "Expected 9,999 archived permutation assignments")
    require(np.array_equal(registry.permutation_id.to_numpy(), np.arange(1, 10000)),
            "Unexpected permutation IDs/order")
    require((registry.seed == 20260615).all(), "Unexpected registry seed")
    require((registry.canonical_order == ",".join(CONDITIONS)).all(), "Unexpected condition coding")
    assignment_values = registry[["assign_A", "assign_B", "assign_C", "assign_P"]].to_numpy()
    require(np.isfinite(assignment_values).all(), "Nonfinite permutation assignment")
    assignments = assignment_values.astype(int)
    require(np.array_equal(assignments, assignment_values), "Permutation assignment is noninteger")
    require(((assignments >= 0) & (assignments < 24)).all(), "Permutation assignment out of range")
    require(len(set(map(tuple, assignments))) == 9999, "Duplicate permutation assignments")
    require(not np.any((assignments == 0).all(axis=1)), "Identity assignment was not excluded")
    permutation_choices = np.array(list(itertools.permutations(range(4))))
    f_permuted = np.empty(9999)
    r2_permuted = np.empty(9999)
    for index, assignment in enumerate(assignments):
        permuted_codes = permutation_choices[assignment[donor_codes], codes]
        # Each donor receives one relabeling for its entire five-day trajectory.
        permuted_ss = residual_ss(full_design(permuted_codes), y)
        permuted_block = reduced_ss - permuted_ss
        f_permuted[index] = (permuted_block / df_block) / (permuted_ss / df_residual)
        r2_permuted[index] = permuted_block / total_ss
    max_f_error = float(np.max(np.abs(f_permuted - registry.F_perm.to_numpy())))
    max_r2_error = float(np.max(np.abs(r2_permuted - registry.delta_R2_perm.to_numpy())))
    require(max_f_error <= ROUNDING_TOLERANCE, "A recomputed permutation F differs beyond rounding")
    require(max_r2_error <= ROUNDING_TOLERANCE, "A recomputed permutation R2 differs beyond rounding")
    extreme = int(np.sum(f_permuted >= observed_f - PERMUTATION_TIE_TOLERANCE))
    p_value = (extreme + 1) / (len(registry) + 1)
    require(extreme == 1, f"Unexpected archived-permutation extreme count: {extreme}")
    require(abs(p_value - float(expected["p_primary"])) < 1e-12,
            "Recomputed primary permutation p differs from archived result")

    report = {
        "status": "PASS",
        "scope": "Primary endpoint only, starting from frozen 80x8 module scores and the archived 9999 assignments",
        "excluded_scope": ["raw-spectrum processing", "protein-level module rescoring", "secondary analyses", "external validation"],
        "package_files_sha256_verified": n_hashes,
        "historical_lf_hashes_verified": n_historical_hashes,
        "input_raw_sha256": digest(input_path),
        "model_reference_coding": "drop-last; independent verifier does not import historical analysis code",
        "n_observations": 80, "n_source_blocks": 4, "n_modules": 8,
        "source_design": "three individual-donor preparations and one preparation pooled from 40 donors",
        "legacy_donor_id": "filename-derived source-block label; P-to-pool mapping inferred, not explicitly verified",
        "biological_independence_verified": False,
        "observed": values,
        "permutation_count": len(registry), "distinct_assignments": 9999,
        "extreme_count": extreme, "permutation_p": p_value,
        "maximum_F_error_vs_six_decimal_registry": max_f_error,
        "maximum_delta_R2_error_vs_six_decimal_registry": max_r2_error,
        "rounding_tolerance": ROUNDING_TOLERANCE,
        "python": platform.python_version(), "numpy": np.__version__, "pandas": pd.__version__,
    }
    output = BASE / "verification/primary_verification.json"
    output.parent.mkdir(exist_ok=True)
    output.write_text(json.dumps(report, indent=2) + "\n", encoding="utf-8")
    print(f"PASS: F={observed_f:.9f}; delta_R2={delta_r2:.9f}; p={p_value:.4f}; b={extreme}; B=9999")
    print(f"Verified {n_hashes} package hashes and every archived primary permutation. Report: {output.name}")
    return report


if __name__ == "__main__":
    try:
        verify()
    except (VerificationError, OSError, ValueError, KeyError) as error:
        print(f"FAIL: {error}", file=sys.stderr)
        sys.exit(1)
```
